# MHC-I binding affinity derived metrics fail to predict tumor specific neoantigen immunogenicity

**DOI:** 10.1101/2022.03.14.484285

**Authors:** Guadalupe Nibeyro, Romina Girotti, Laura Prato, Gabriel Moron, Hugo D. Luján, Elmer A. Fernandez

## Abstract

Tumor-specific antigens emerging through somatic genomic rearrangements, known as neoantigens, play a critical role in current anticancer immunotherapy. They may or may not elicit an immune response when presented on the tumor cell surface bound to the MHC-I molecule, whose strength has been assumed as an indicator of immunogenicity. Several *in silico* peptide-MHC-I binding affinity predictors are used to prioritize putative immunogenic neoantigens to be experimentally and clinically explored either as biomarkers or targets for anticancer vaccines. This claims for a fair evaluation of such predictors, making essential the development of appropriate databases with experimentally validated, immunogenic/non-immunogenic neoantigens. Thus far, such a database is lacking. We herein present ITSNdb, a new and curated immunogenic neoantigen database and use it to benchmark current neoantigen immunogenicity predictors. Benchmark results failed to support the application of the predicted peptide-MHC-I binding affinity or its derived metrics as a tool to estimate neoantigen immunogenicity and the tumor neoantigen burden as an immunotherapy response biomarker. Moreover, binding affinity based immunogenicity definition leads to identifying wild-type peptide counterparts as predictors of immunotherapy response. We demonstrate that MHC-I binding affinity is insufficient to define neoantigen immunogenicity, despite being necessary for neoantigen tumor cell presentation suggesting that a paradigm shift for the emergence of new rules to identify immunogenic neoantigens is required.

## Introduction

Current immunotherapy has become a mature cancer treatment strategy in addition to surgery, chemotherapy, and radiotherapy; showing significant therapeutic effects in many human tumors by harboring the immune system’s capacity to eliminate cancer cells [1]. However, a substantial number of patients do not benefit from this treatment approach [2]. Current checkpoint blockade immunotherapy (ICB) targeting PD1, PD-L1 and CTLA4 takes advantage of immune cell infiltration in the tumor to reinvigorate an efficacious antitumoral immune response. Neoantigen-specific T cells are critical for effective ICB.

Briefly, neoantigens are unique antigenic peptides that emerge from genomic alterations encompassing single nucleotide variants (SNVs), nucleotide insertions or deletions, alternative splicing and/or gene fusion events [3-6], among others. These alterations may produce dysfunctional proteins by either non-synonymous mutations or changes to the open reading frames [3,4]. When processed by the proteasome, different neopeptides can be generated, presented on the cell surface bound to the MHC-I molecule (i.e., neoantigens) and, potentially, trigger an immune response if recognized by T cell receptors [7]. Current development of alternative therapeutic strategies, like adoptive T cell transfer [8] and neoantigens based vaccines [9], are grounded on the existence of immunogenic neoantigens in several tumor types.

The advent of sequencing technologies has allowed for genome-wide and transcriptome-wide scanning of potential neoantigens on an unprecedented scale, impacting patient care. For instance, high tumor mutation burden (H-TMB) or its surrogate, the tumor neoantigen burden (TNB) [1,7] (only including neopeptides predicted to bind to the MHC-I) are leading candidate biomarkers for identifying cancer patients who may benefit from ICB. These biomarkers are based on the underlying assumption that high numbers of altered proteins imply high numbers of antigenic peptides; thus, increasing anti-tumor immunogenicity [2]. However, poorly mutated tumors also benefit from the appearance of immunogenic neoantigens derived from other types of genomic rearrangements and non-synonymous SNVs [10].

Based on these findings, several *in silico* tools were developed to predict immunogenic neoantigens from genomic- or transcriptomic-derived sequence data [11-13]. Most of them assume that the capability of a neopeptide to be presented on the cell surface, strongly attached to some MHC-I molecule, is sufficient to elicit an immune response directing the use of binding affinity (BA) predictors over HLA-restricted detected neopeptides. Thus, different BA and derived score thresholds are currently used to define neoantigen immunogenicity broadly used in most genomic-based neoantigen search tools, despite that these thresholds were settled on viral or non-self-antigen peptides [11-13].

Although BA predictors have been well validated as such, their use to predict immunogenic neoantigens is highly controversial since only a small fraction have been experimentally validated as being positive immunogenic [14]. This implies that these tools, designed to predict affinity between the peptide and the MHC-I molecule, may provide high false positive rates when used to define putative immunogenic neoantigens, making their use both impractical and financially ineffective for personalized vaccine development. Moreover, since immunogenic neoantigens are patient-specific, neoantigen immunogenicity predictors should not only be sensitive and specific but also highly confident (i.e., high positive and negative predictability). Therefore, to evaluate predictor’s performance, databases with truly validated immunogenic and non-immunogenic tumor derived neoantigens become essential.

Currently used neoantigen immunogenic prediction tools and/or metrics were developed using viral and/or non-self-peptides from the <u>I</u>mmune <u>E</u>pitope <u>D</u>ata<u>b</u>ase (IEDB) [15] and evaluated using few validated immunogenic neoantigens (positive), and natural human peptides (negative) immunogenic examples [16]. However, it has been reported that peptides derived from natural proteins may also bind to the MHC-I molecule and consequently be present on the cell surface [17]. Therefore, the selection or prioritization of peptides as potentially immunogenic based solely on BA scores or derived metrics may result in false positive predictions (i.e., truly negative examples predicted as immunogenic). Moreover, the lack of experimentally validated non-immunogenic tumor-derived neoantigens hinders the fair performance evaluation of the current methods. In this regard, several attempts have been conducted to develop tumor specific antigens (TSAs) databases, such as the <u>C</u>ancer <u>A</u>ntigenic <u>P</u>eptide <u>D</u>atabase (CAPD) [18] and *DbPepNeo* [19], but they lack truly non-immunogenic peptides, limiting the evaluation of false positive rates. The recently developed TANTIGEN database [20], incorporates tumor T cell antigens, but information regarding experimental MHC-I binding, validation method or immunogenicity classification is not clearly accessible. Other databases containing tumor immunopeptidomics and incorporating neoantigens have emerged, such as *caAtlas* [21], but they lack information about peptide immunogenicity, making it impractical for predictor performance assessment.

Here, to yield a fair comparison of tumor immunogenic neoantigen predictors, a revision and curation of current databases and literature has been performed, providing the first <u>I</u>mmunogenic <u>T</u>umor-<u>S</u>pecific <u>N</u>eoantigen <u>d</u>ata<u>b</u>ase (ITSNdb) holding a reliable list of well-described tumor-derived neoantigens with reported experimental validation of their MHC-I binding and their positive or negative immunogenic reaction (i.e. immunogenic and non-immunogenic neoantigens, respectively). Through this new database, a comprehensive immunogenic predictor’s assessment was performed, showing the deficiencies of current methods and metrics to predict or prioritize immunogenic tumor neoantigens.

## Methods and data

### Database constitution

A manually curated neoantigen database was created using a novel approach. The ITSNdb only includes neoantigens meeting the following inclusion criteria: (a) peptides derived from non-silent somatic SNVs; (b) with experimentally validated binding to the MHC-I molecule; (c) tested on immunogenic assays (i.e., tetramer titration and IFN-γ or TNF ELISPOT^®^); and (d) their wild-type (WT) sequence, present in referenced protein sequence. All the ITSNdb peptides possess experimentally-validated positive or negative immunogenic activity thus being classified as “Positive” or “Negative” neoantigens, respectively (see supplementary material for more details).

Up to now, the ITSNdb, holds 61 nine-mer SNV-derived neoantigens with their: WT counterparts, restricted HLA information, gene, tumor tissue and reference (Supplementary table 1). Figure 1A describes the current peptide distribution among tumor types, as well as their immunogenicity, mutation position and mutation position type (i.e., MHC-I anchor position if the amino acid change is located at position 2 or 9 in the sequence, or MHC-I non-anchor position). Figure 1B shows the peptide count distribution for HLA restriction type. The HLA A02:01 represents 39/61 (63.93%) of the HLA restricted peptides.

**Fig 1.**
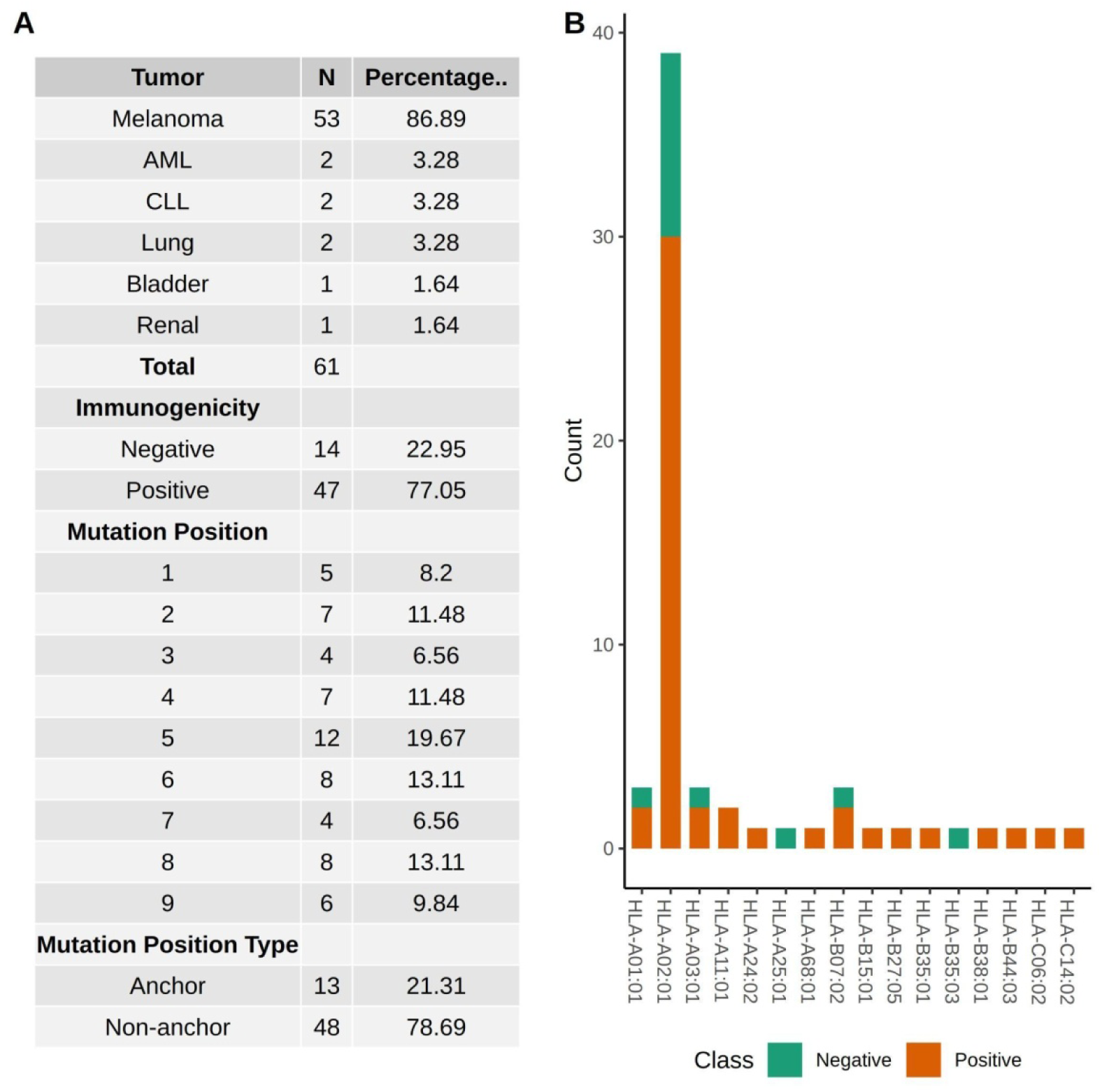
Database composition. (A) Detailed description of the characteristics of neoantigens included in the current database. (B) Distribution of positive (immunogenic) and negative (non-immunogenic) neoantigens by HLA subtype.

### Methods

To evaluate the performance of current methods and metrics to assess neoantigen immunogenicity, seven state-of-the-art types of software were run using the authors’ default preferences over ITSNdb. The evaluated software could be divided into two different categories, those predicting BA scores: netMHCpan 4.1 [22], MHCflurry 2.0 [23], and mixMHCpred 2.1 [24]; and those predicting immunogenicity scores or classes: deepimmune-CNN 1.2 [25], T cell class I pMHC immunogenicity predictor (CIImm) [26], deepitope and PRIME 1.0 [27] (see supplementary material for more information). In addition, the <u>D</u>ifferential <u>A</u>gretopicity <u>I</u>ndex (DAI), derived from the predicted BA metrics, was also evaluated. DAI was originally proposed as the BA difference between WT and its mutated counterpart predicted by netMHCpan_BA_ (I- DAI_BA_=WT_BA_-mut_BA_) [28], where higher DAI is interpreted as a higher chance of immunogenicity, under the assumption that peptides with amino acid substitutions should have a stronger affinity to MHC-I to trigger an immune response. Later the ratios between predicted percentage ranks (II-DAI_%R_=mut_%R_/WT_%R_) [29] or BA (II-DAI_BA_) [30] were proposed respectively, where lower DAI score is interpreted as a higher chance of immunogenicity. There is no consensus about its calculation procedure, but the concept remains the same under the assumption that the DAI score correlates with neoantigen immunogenicity. Here, I-DAI and II-DAI were calculated with all the BA and ranks predicted by both netMHCpan (x-DAI_BA_, x-DAI_%R_) and MHCflurry (x-DAI_BA_, x-DAI_P_), where x=I,II.

### Statistical analysis

All the predicted scores from all the considered methods and their derived metrics were evaluated through ROC curves (pROC R library [31]) and the area under the curve (AUC). For evaluated metrics, the optimal classification threshold was estimated using the methods provided by the pROC library (i.e., Youden’s method, see equation 1) and by the Distance to the Optimal Point [32] (DOP, see equation 2). For the established optimal threshold, the sensitivity (Se), specificity (Sp), positive and negative predictability values (PPV and NPV), F1 score for positive predictions and false positive rates (FPR) for neoantigen immunogenicity classification were used for comparisons. These procedures were also used to evaluate predictions onto WT counterpart peptides.

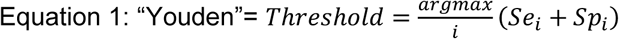

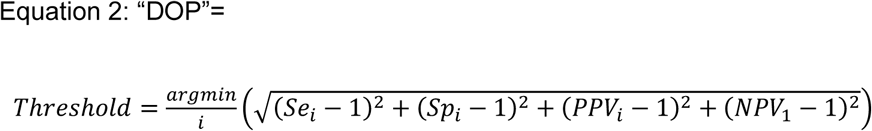

### Validation peptides

#### Negative set

109 neopeptides with experimental negative immunogenic assay (4 with 2 HLA associations) of 9-mer length derived from SNVs (most of them originally predicted as binders by netMHCpan version 4.0; thus, putatively predicted as immunogenic), were collected from Robbins *et al*. [3]. They were not included in the ITSNdb due to missing binding affinity experimental validation. (Supplementary Table 2).

#### Positive set

Six non-SNV derived neoantigens of 9-mer length, with validated MHC-I presentation and immunogenicity [4-6] were used. Two neoantigens derived from gene fusion events; one of them with 2 HLA-associated restriction alleles. Three originated from intronic retention. The last one was derived from 2 nucleotide deletions resulting in a new open reading frame (Supplementary Table 3). The absence of the WT counterparts was the reason for not including them in ITSNdb.

### Immunotherapy response clinical trials datasets

Three ICB treated cohorts (one non–small cell lung and two melanoma cancer cohorts) [33-35] were used to evaluate the impact of immunogenic prediction on TNB and their WT peptide counterparts (See supplementary material for more information).

### Vaccine design neoantigen prioritization datasets

Three prioritized UVB-derived neoantigens from melanoma mice tumors [36] and two prioritized mRNA vaccine-induced neoantigens for gastrointestinal cancer clinical trial [37] were evaluated (supplementary table 4)

## Results

### ROC performance metrics were similar using mutated or wild type peptides

Figure 2 shows the ROC plots for each evaluated method and score (except for Deepitope) applied to mutated and WT peptides (Figures 2A and 2B, respectively). The DAI scores are shown in Figure 2C. The netMHCpan_BA_ reached the highest AUC (0.687); followed by DeepImmune (0.682) and MHCflurry_BA_ (0.629) (Table 1). The AUCs’ DAI scores ranged from 0.457 to 0.568, resembling a random classification. Noteworthy, when fed by WT, AUCs of DeepImmune (AUC=0.693), mixMHCpred_R_ (AUC=0.549) and mixMCHpred_S_ (AUC=0.543) were higher than when fed by their mutated counterpart peptides.

**Table 1.**
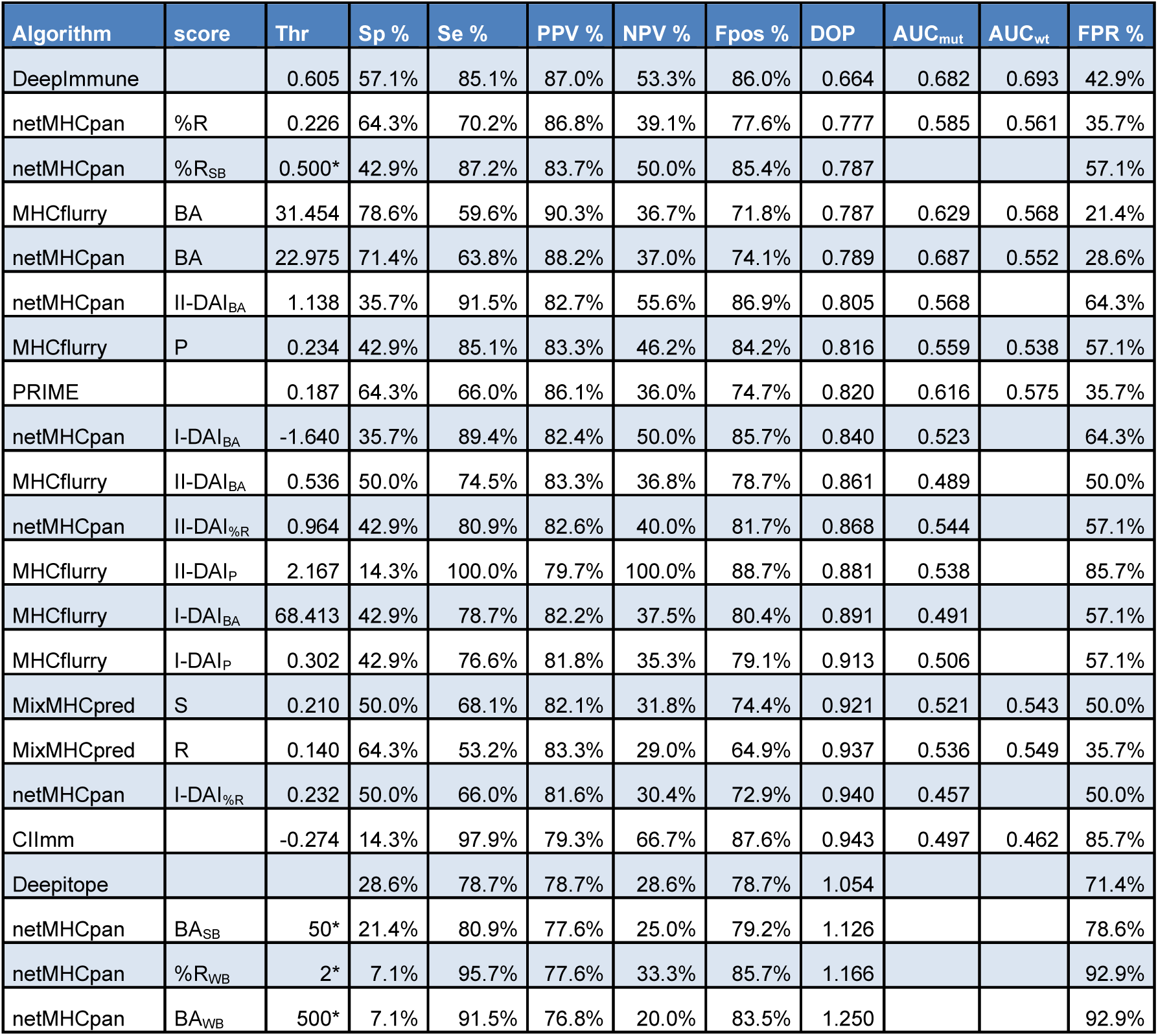
Performance values calculated for each metric. Thr*: authors’ proposed threshold. I-DAI: WT-mut. II-DAI: mut/WT.

| Algorithm | score | Thr | Sp % | Se % | PPV % | NPV % | Fpos % | DOP | AUC<sub>mut</sub> | AUC<sub>wt</sub> | FPR % |
| --- | --- | --- | --- | --- | --- | --- | --- | --- | --- | --- | --- |
| DeepImmune |  | 0.605 | 57.1% | 85.1% | 87.0% | 53.3% | 86.0% | 0.664 | 0.682 | 0.693 | 42.9% |
| netMHCpan | %R | 0.226 | 64.3% | 70.2% | 86.8% | 39.1% | 77.6% | 0.777 | 0.585 | 0.561 | 35.7% |
| netMHCpan | %R<sub>SB</sub> | 0.500* | 42.9% | 87.2% | 83.7% | 50.0% | 85.4% | 0.787 |  |  | 57.1% |
| MHCflurry | BA | 31.454 | 78.6% | 59.6% | 90.3% | 36.7% | 71.8% | 0.787 | 0.629 | 0.568 | 21.4% |
| netMHCpan | BA | 22.975 | 71.4% | 63.8% | 88.2% | 37.0% | 74.1% | 0.789 | 0.687 | 0.552 | 28.6% |
| netMHCpan | II-DAI<sub>BA</sub> | 1.138 | 35.7% | 91.5% | 82.7% | 55.6% | 86.9% | 0.805 | 0.568 |  | 64.3% |
| MHCflurry | P | 0.234 | 42.9% | 85.1% | 83.3% | 46.2% | 84.2% | 0.816 | 0.559 | 0.538 | 57.1% |
| PRIME |  | 0.187 | 64.3% | 66.0% | 86.1% | 36.0% | 74.7% | 0.820 | 0.616 | 0.575 | 35.7% |
| netMHCpan | I-DAI<sub>BA</sub> | -1.640 | 35.7% | 89.4% | 82.4% | 50.0% | 85.7% | 0.840 | 0.523 |  | 64.3% |
| MHCflurry | II-DAI<sub>BA</sub> | 0.536 | 50.0% | 74.5% | 83.3% | 36.8% | 78.7% | 0.861 | 0.489 |  | 50.0% |
| netMHCpan | II-DAI<sub>%R</sub> | 0.964 | 42.9% | 80.9% | 82.6% | 40.0% | 81.7% | 0.868 | 0.544 |  | 57.1% |
| MHCflurry | II-DAI<sub>P</sub> | 2.167 | 14.3% | 100.0% | 79.7% | 100.0% | 88.7% | 0.881 | 0.538 |  | 85.7% |
| MHCflurry | I-DAI<sub>BA</sub> | 68.413 | 42.9% | 78.7% | 82.2% | 37.5% | 80.4% | 0.891 | 0.491 |  | 57.1% |
| MHCflurry | I-DAI<sub>P</sub> | 0.302 | 42.9% | 76.6% | 81.8% | 35.3% | 79.1% | 0.913 | 0.506 |  | 57.1% |
| MixMHCpred | S | 0.210 | 50.0% | 68.1% | 82.1% | 31.8% | 74.4% | 0.921 | 0.521 | 0.543 | 50.0% |
| MixMHCpred | R | 0.140 | 64.3% | 53.2% | 83.3% | 29.0% | 64.9% | 0.937 | 0.536 | 0.549 | 35.7% |
| netMHCpan | I-DAI<sub>%R</sub> | 0.232 | 50.0% | 66.0% | 81.6% | 30.4% | 72.9% | 0.940 | 0.457 |  | 50.0% |
| CIImm |  | -0.274 | 14.3% | 97.9% | 79.3% | 66.7% | 87.6% | 0.943 | 0.497 | 0.462 | 85.7% |
| Deepitope |  |  | 28.6% | 78.7% | 78.7% | 28.6% | 78.7% | 1.054 |  |  | 71.4% |
| netMHCpan | BA<sub>SB</sub> | 50* | 21.4% | 80.9% | 77.6% | 25.0% | 79.2% | 1.126 |  |  | 78.6% |
| netMHCpan | %R<sub>WB</sub> | 2* | 7.1% | 95.7% | 77.6% | 33.3% | 85.7% | 1.166 |  |  | 92.9% |
| netMHCpan | BA<sub>WB</sub> | 500* | 7.1% | 91.5% | 76.8% | 20.0% | 83.5% | 1.250 |  |  | 92.9% |

**Fig 2.**
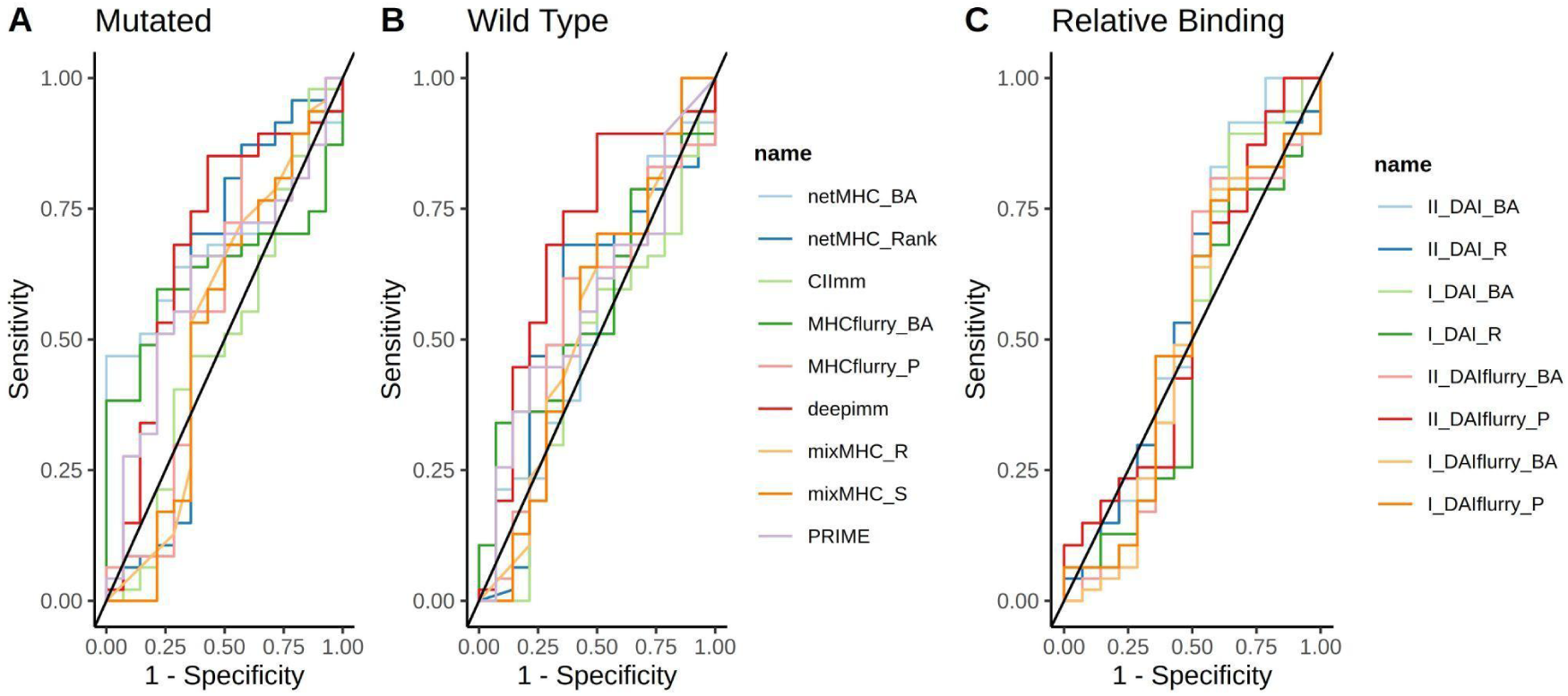
ROC curves for different softwares and direct metrics: mutated peptides (A) and wild type counterparts (B) are shown. ROC curves for derived metric: all calculated DAI (C).

### The Distance to the Optimal Point provides better performance metrics

To evaluate the different methods for neoantigen immunogenicity predictions, it was necessary to choose a classification threshold over the continuous predicted scores. Figure 3 shows the comparison between F1 positive score (Figure 3A) and *Se* (Figure 3B) achieved by the threshold calculated through Youden and DOP methods (Equations 1 and 2, respectively). By using DOP, the resulting performance metrics were superior (Wilcoxon paired test p<0.05). Thus, the performance metrics presented in Table 1 were established using the classification score threshold derived by DOP method.

**Fig 3.**
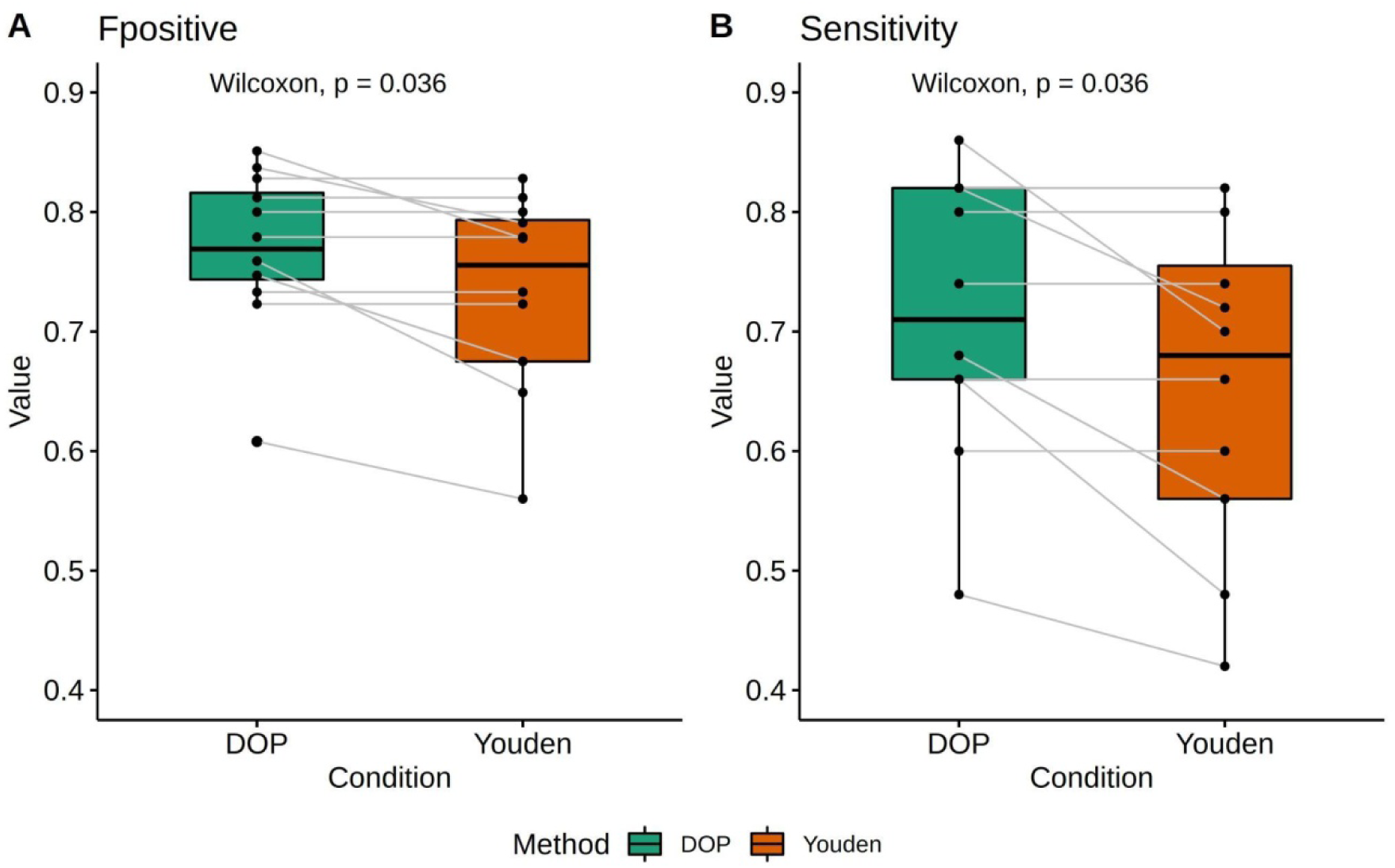
Comparison of DOP and Youden methods to calculate thresholds according to Fpositive values (A), and Sensitivity values (B).

### None of the evaluated methods performed better than the others

Table 1 shows the performance metrics (*Se, Sp, PPV, NPV, FPR, F1* positive scores and *DOP*) for each method and derived metrics obtained by using the classification thresholds selected with the *DOP* (Equation 2) and with those suggested by the authors’ methods. Sorted by *DOP*, which accounts for both detection performance and certainty, DeepImmune ranked first followed by netMHCpan_%R_, netMHCpan_%R_SB_ and MHCflurry_BA_. However, MHCflurry_BA_ reached the highest *Sp* and PPV but with lower *Se* and NPV than the other three software, suggestive of a higher false negative rate.

For DAI scores, the Sp was around 50% in all cases, with an AUC not significantly different from 0.5 (Wilcoxon test p=0.889), resembling a random classification. The II-DAI_P_ (based on MHCflurry) achieved the highest NPV. It reached a *Se* of 100% (no false negatives) but at the expense of low non-immunogenic detection rate (Sp=14.2%), indicating that almost all neoantigens were predicted as immunogenic (high FPR).

The netMHCpan_BA_ (chosen by DOP) reached a *Se* of 64% meaning that approximately one in 3 immunogenic neoantigens would be missed with this method and about 1 in 4 positive peptides would be misclassified (FPR 28.6%). DeepImmune reached higher Se (85%) with lower *Sp* (57%, i.e many of the negative neoantigens would represent false positive discoveries). The highest *Se* for detecting positive immunogenic peptides was achieved by the II-DAI_P_ derived from MHCflurry, followed by CIImm and the classification based on the WB score threshold for %R=2% proposed by the netMHCpan’s authors. However, they were accomplished at the expense of having low *Sp* values (14.3%, 14.3% and 7,1% respectively). These results imply high false positive rates, and the inability to recognize negative immunogenic peptides. The highest *Sp* (78.6%) was achieved by using the DOP-based threshold of 31.45 nM from MHCflurry_BA_, but at low confidence levels (NPV≃ 36.7%) and a positive detection rate (*Se*) of 59.6%.

From the methods that predicted direct immunogenicity scores or proposed direct classification (DeepImmune, CIImm, Deepitope and PRIME), CIImm reached the highest immunogenic detection rate (Se=97.9%) but with a FPR of 85.7% meanwhile DeepImmune provided a Se=85.1% with an FPR of 42.9%. Deepitope reached a high Se=80% with an FPR of 62.5% and PRIME with a *Se* of 66% and an *Sp* of 64.3%.

Although the sensitivity of all tested methods was acceptable, the FPR varied greatly, ranging from ∼93% (netMHCpan_BA_ WB-500 nM and netMHCpan_%R_ WB-2%) to ∼21% for MHCflurry_BA_. This observation is consistent with previous reports where only a very low fraction of predicted immunogenic neoantigens were experimentally confirmed (high false positives rates) resulting in a waste of resources and negatively impacting their widespread use as predictors of neoantigen immunogenicity.

### DAI scores are indicators of position type mutation but not of immunogenicity

The Differential Agretopicity Index (DAI) suggests that the relative BA between mutated and WT peptides is an indicator of immunogenicity under the hypothesis that mutated peptides have a stronger MHC-I affinity than their WT counterpart [28]. Figure 4 shows the distribution of the DAI (I-DAI_BA_=WT_BA_-mut_BA_; where higher values imply higher chances of being immunogenic) derived from netMHCpan and MHCflurry BA predictions (For other DAI see supplementary material). Significant differences were found (wilcoxon p<0.01) comparing if the mutation position lies in an anchor or non-anchor position (Figure 4A) but no significant differences between positive and negative immunogenicity (Figure 4B) were observed. These results, in accordance with Capietto *et al*. [29], suggest that, since the amino acids located at anchor positions 2 and 9 (C-terminal) are responsible for peptide-MHC-I binding, predicted BA will be more affected by mutations in such positions than those located in non-anchor. On the other hand, regarding immunogenic vs. non-immunogenic DAI-based predictions, none of the DAI scores could differentiate between such classes, as shown by the ROC analysis (Figure 2C and Table 1). These findings suggest that the DAI is indicative of whether the mutation occurs at an anchor position or not, but doesn’t provide information regarding the immunogenic capability of the evaluated peptide.

**Fig 4.**
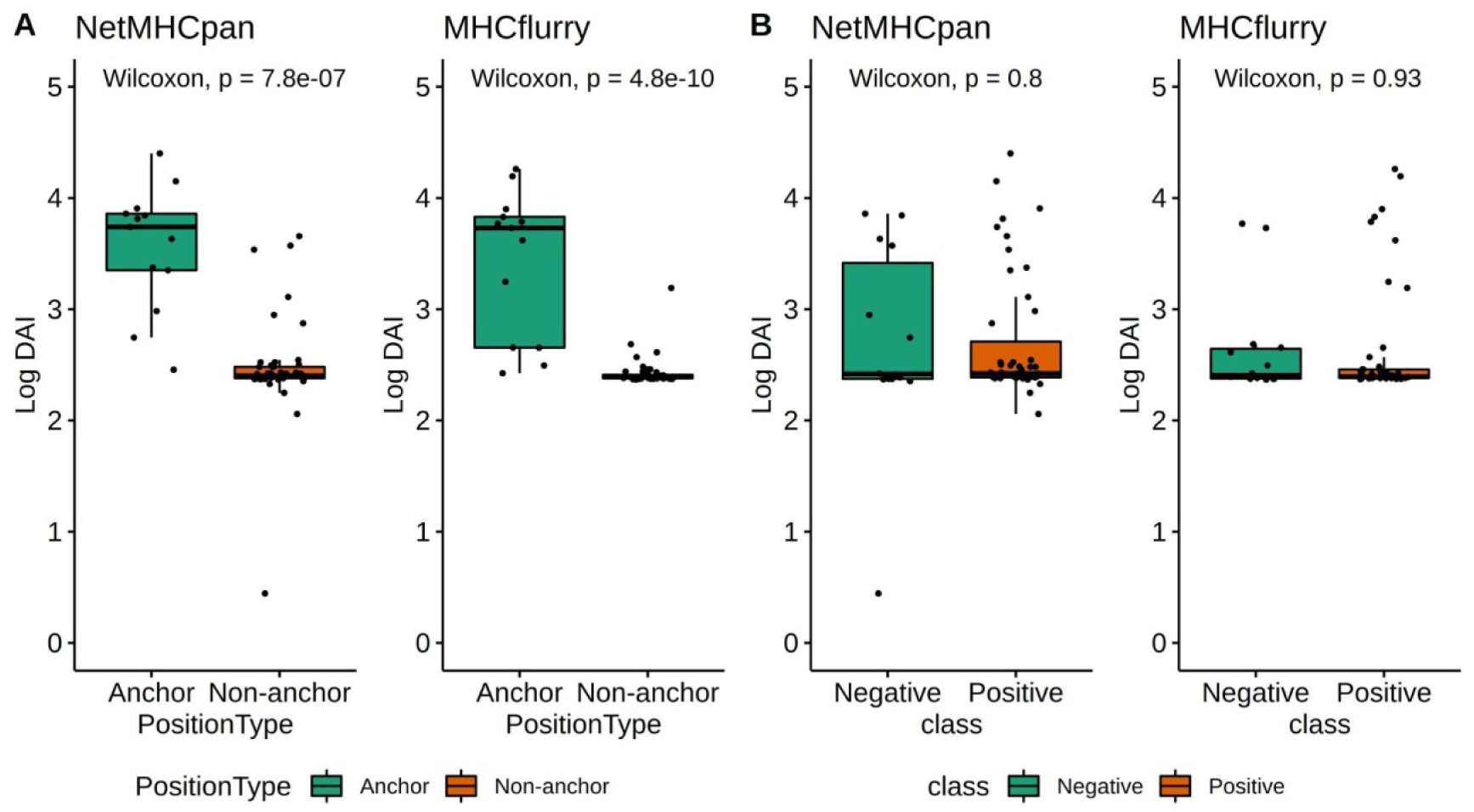
(A) Comparison between DAI values for netMHCpan and MHCflurry wild-type and mutated peptides BA difference and (B) according to Position Type (anchor vs non-anchor) and Class (negative vs positive). DAI were calculated as log(DAI + k) for visualization purposes.

### Wild-type counterpart peptides were also classified as immunogenic in both immunogenic and non-immunogenic neoantigen classes

It’s been reported that the WT counterpart of neoantigens can be presented by the MHC-I molecule and they may elicit an initial immune response that, in mice, was then self-limited [28,38] and in humans[16] demonstrated immunogenicity at high concentrations [3]. This is in accordance with the exposed results that, for some metrics (DeepImmune and mixMHCpred), the AUCs were higher for WT than for their mutated counterpart and the low DAI values obtained for immunogenic neoantigens with non-anchor mutations. Evaluating immunogenicity predictions for the WT counterparts based on Table 1 thresholds, Figure 5 shows the amount of peptide pairs where the mut/WT peptide was classified as immunogenic (I) or as non-immunogenic (NI) for both immunogenic and non immunogenic classes (Figure 5A and 5B respectively).

**Fig 5.**
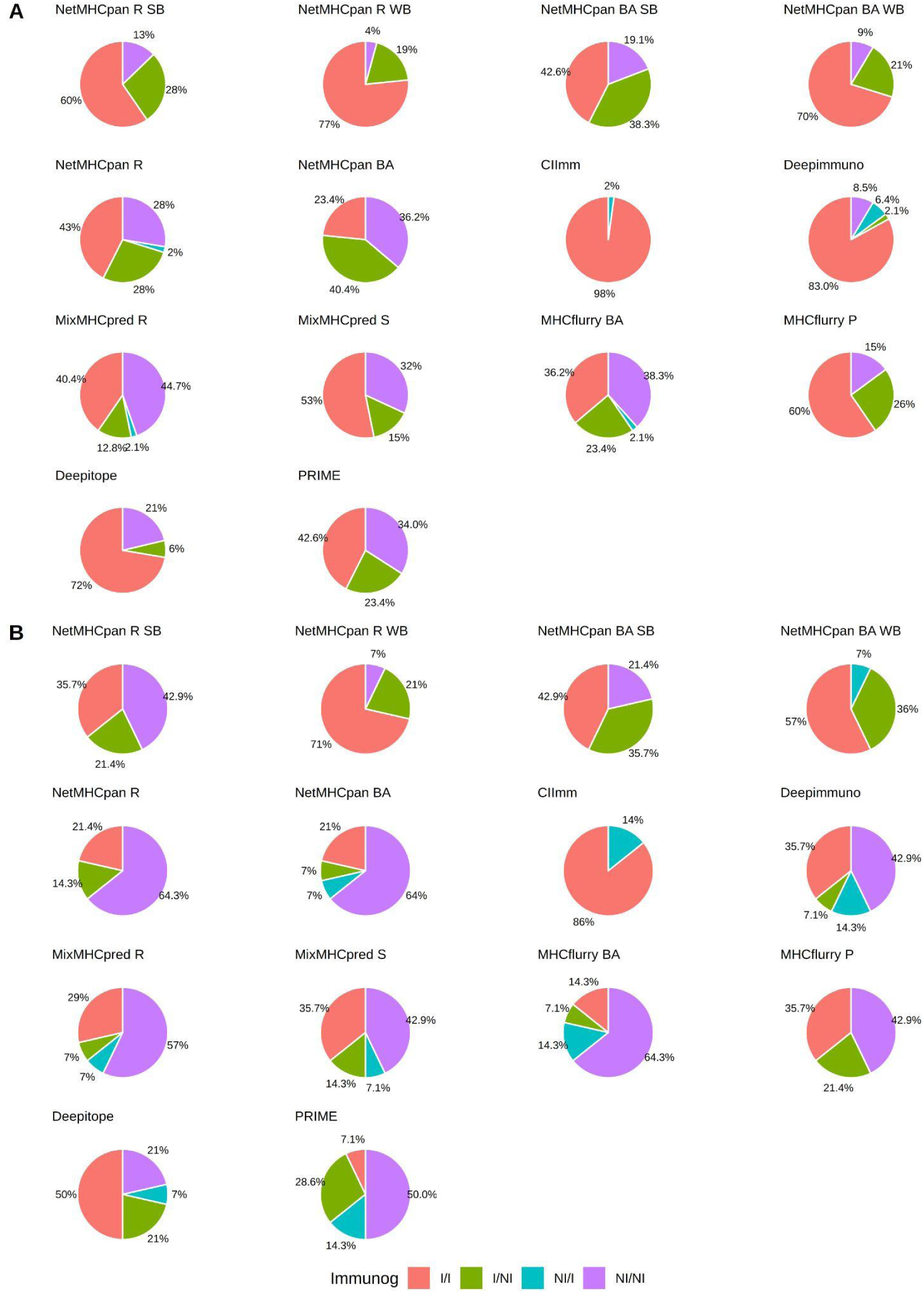
Mutated and WT counterpart neoantigen predicted classification into immunogenic (I) or non-immunogenic (NI), according to different metrics, using DOP and authors thresholds. (A) Positive neoantigens. (B) Negative neoantigens.

For immunogenic neoantigens (Figure 5A), 60% and 77% of the WT counterpart peptides were classified as immunogenic using netMHCpan_%R_ with SB and WB classification thresholds respectively; and 42.6% and 70% when using netMHCpan_BA_ (SB and WB, respectively). Using netMHCpan_BA_ with SB definition, 38.3% of the WT peptides were recognised as non-immunogenic and 19.1% of the mut/WT peptides pairs were both classified as non-immunogenic.

The higher proportion of peptide pairs where the mutated peptide was classified as immunogenic and their WT counterpart as non-immunogenic (identified as I/NI), was achieved by netMHCpan_BA_ using the DOP-based threshold (40.4%). CIImm achieved the highest proportion of immunogenic classification for both mutated and WT pairs (98%) (I/I). MixMHCpred_R_ classified the highest proportion (44.7%) of both peptides (mut/WT) as non-immunogenic (NI/NI) and Deepimmuno achieved the highest proportion (6.4%) of peptide pairs by which the mutated peptide was classified as non-immunogenic and the WT as immunogenic (NI/I).

For non-immunogenic neoantigens (Figure 5B), the DOP-based threshold improved the non-immunogenic classification on both peptides of the mut/WT pair reaching 64.3% and 64% for netMHCpan_%R_ and netMHCpan_BA_ respectively, compared to using SB (42.9% and 21.4%) and WB (7% and 0%) thresholds. These results were similar for MHCflurry_BA_. Curiously, CIImm predicted all WT as immunogenic.

### Current methods and classifiers based on BA metrics yield high false positive rates on non-immunogenic neopeptide sets

The 109 non-immunogenic neopeptides from the negative validation set (four of them had 2 HLA associated restrictions; thus considered as 2 different peptides, 113 in total, for those HLA dependent predictors) were used to evaluate FPR of prediction software. All these neopeptides were originally classified as putatively immunogenic by netMHCpan version 4.0[3]

Table 2 shows the amount of neopeptides predicted as negative (non-immunogenic) according to DOP, authors’ thresholds, and by the classification tag. The highest performance was achieved by netMHCpan_BA_ (version 4.1) predicting 88 neopeptides over 113 as negatives, thus yielding an FPR of 22.12%, followed by MHCflurry_BA_ and DeepImmune with a FPR of 24.78 and 26.55% respectively. These were in agreement with Table 1, where netMHCpan_BA_ displayed the second highest specificity with a low FPR (28.57%). The second-best performance was achieved by MHCflurry_BA_, which showed the highest Sp. The SB-based NetMHCpan_%R_, showed the poorest performance, classifying only 5 neopeptides as negative. Interestingly, this method is currently the most widely used for neoantigen immunogenicity prediction, and classified only 28 neopeptides as negative, thus misclassifing 75.22% of the negative neoantigens as positives and the newest netMHCpan 4.1 version improves the FPR compared with the previous version 4.0 and even more if using the DOP-based classification threshold. Largely, the FPR ranges between 22.12% (netMHCpan_BA_) to 95.58% (netMHCpan_%R_ WB) with a median value of 56.03%.

**Table 2.**
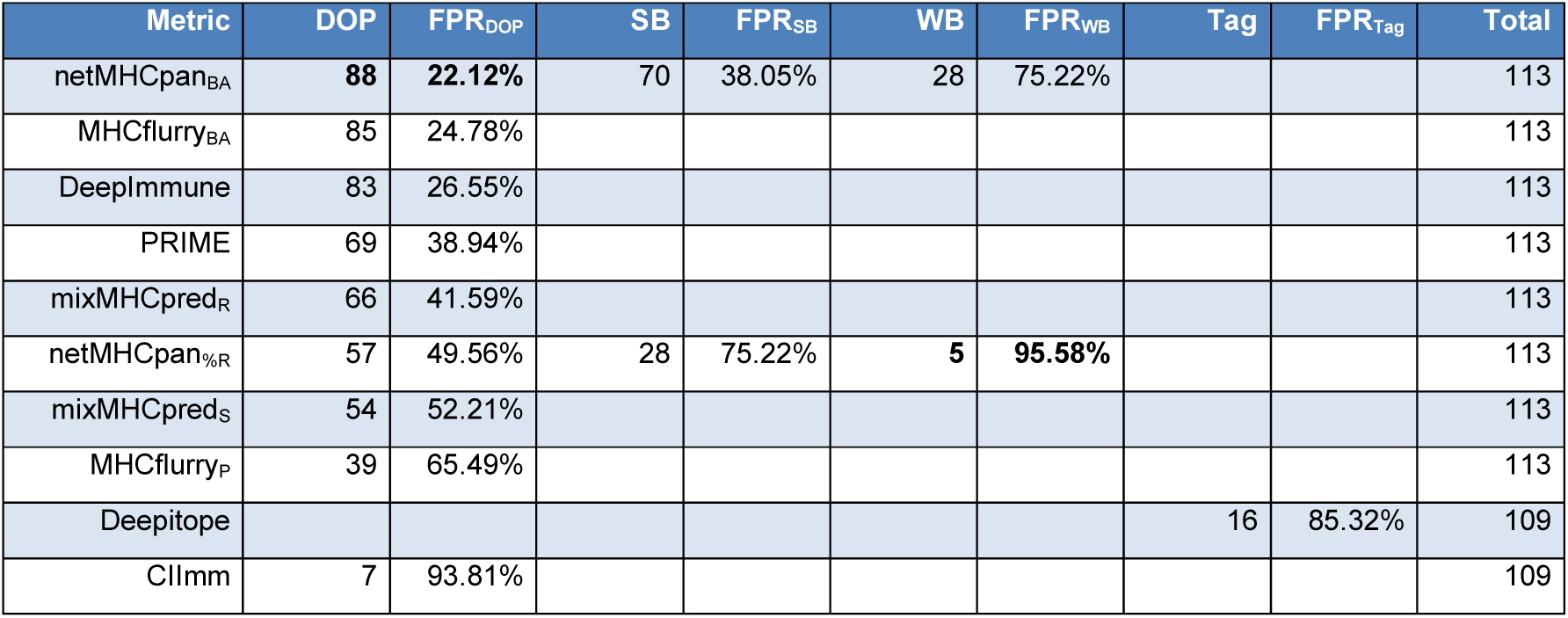
Non-immunogenic peptides predicted as negatives. DOP: distance to the optimal point. FPR: false positive rate. SB: strong binder. WB: weak binder.

### Current BA-based methods achieved high false negative rates over non-point mutation derived immunogenic neoantigens

Immunogenic neoantigens not only derive from point mutations, but also from other genomic events. Here, six experimentally validated immunogenic neoantigens derived from intronic retention events [5], gene fusion events [6] and new open reading frames due to a 2 bp deletion [4] were evaluated using the different thresholds presented in Table 1. One of them had 2 associated HLA, thus it was evaluated as two different peptide-MHC-I complexes when required by the software.

Table 3 shows the number of peptides predicted as immunogenic by each method using DOP, authors’ thresholds or by classification tag (Deepitope). Only CIImm could identify all the neoantigens as immunogenic. Among the others, the top performer was DeepImmune, predicting 6 out of 7 peptides as immunogenic, reaching a FNR of 14.29% in accordance with Table 1. Interestingly, netMHCpan_BA_ recognized 1 out of 7 peptides as immunogenic when using the DOP-based classification threshold and 3 and 4 out of 7 when using SB and WB thresholds respectively. This algorithm increases Sensitivity at the expense of lower certainty (lower positive predictability) as stated in Table 1. In general, the FNR ranges from 0% (CIImm) to 85.71% (netMHCpan_BA_ DOP), with a median value of 41.84%.

**Table 3.**
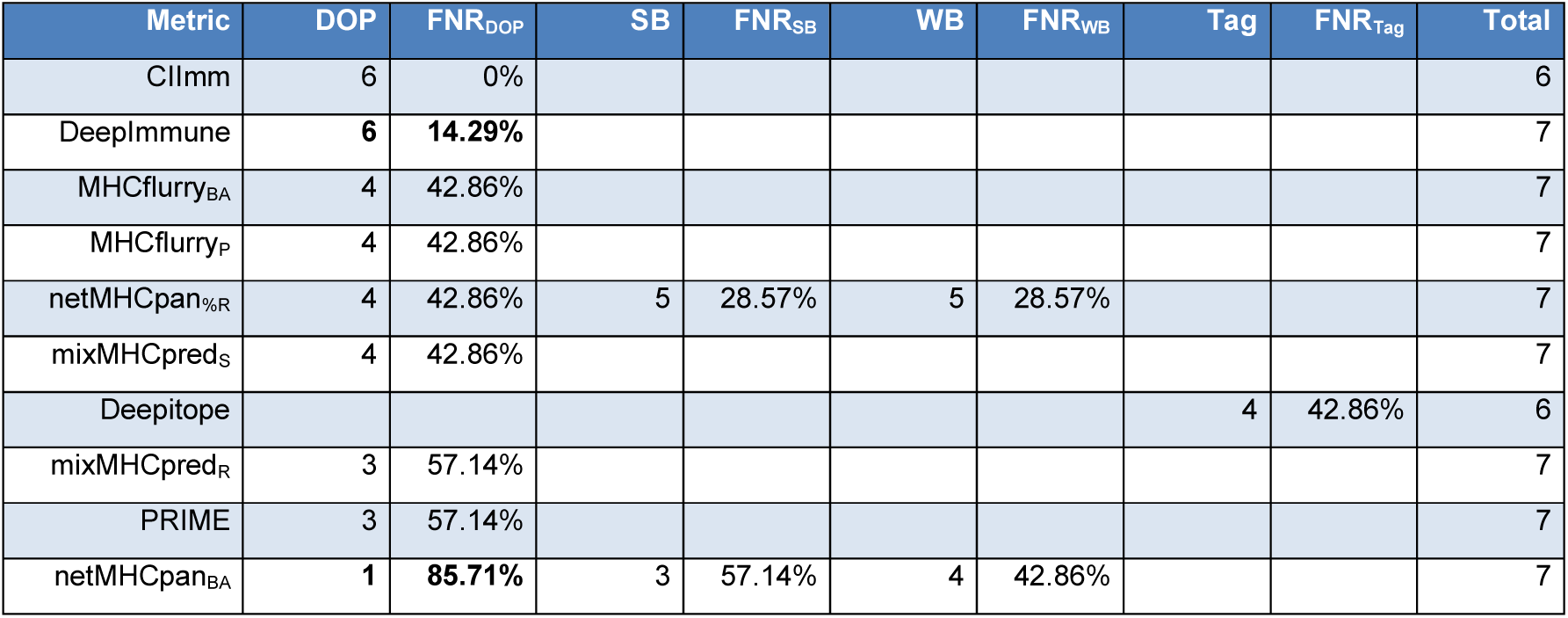
Immunogenic peptides predicted as positives. DOP: distance to the optimal point. FPR: false positive rate. SB: strong binder. WB: weak binder.

### Binding affinity scores predicted wild-type peptides as immunogenic with ICB response association

Tumor Neoantigen Burden (TNB) has been suggested as an ICB biomarker like the Tumor Mutation Burden (TMB) [1,7,33-35]. The TNB is derived from the TMB by counting those neoantigens predicted to be binded to the MHC-I molecules by BA predictors. Based on the shown immunogenic prediction on WT peptides, neoantigens and their WT derived peptides from three ICB clinical trials were evaluated.

Exploring *Rizvi et al*. [33] dataset, in accordance to the original publication, the TNB, calculated from neoantigens derived from non-synonymous mutations with predicted BA<500nM to the MHC-I molecule (with netMHC version 3.4) [39], was found to be associated to durable clinical benefit (DCB) beyond six months (Wilcoxon test p=0.0018; figure 6A, left panel). However, the TNB calculated using the number of WT counterpart peptides predicted to be binded to the MHC-I molecule, was also found to be associated with DCB (Wilcoxon test p=0.0037; Figure 6A, right panel). Additionally, the WT counterpart ASNA**P**SAAK of the immunogenic neoantigen ASNA**S**SAAK (validated by *Rizvi et al*.), reached a lower BA and a lower %R (BA_WT_=10.39 and %R_WT_=0.033) than the mutated one (BA_mut_=14.47 and %R_mut_=0.069). Thus, being less immunogenic according to the DAI concept.

**Fig 6.**
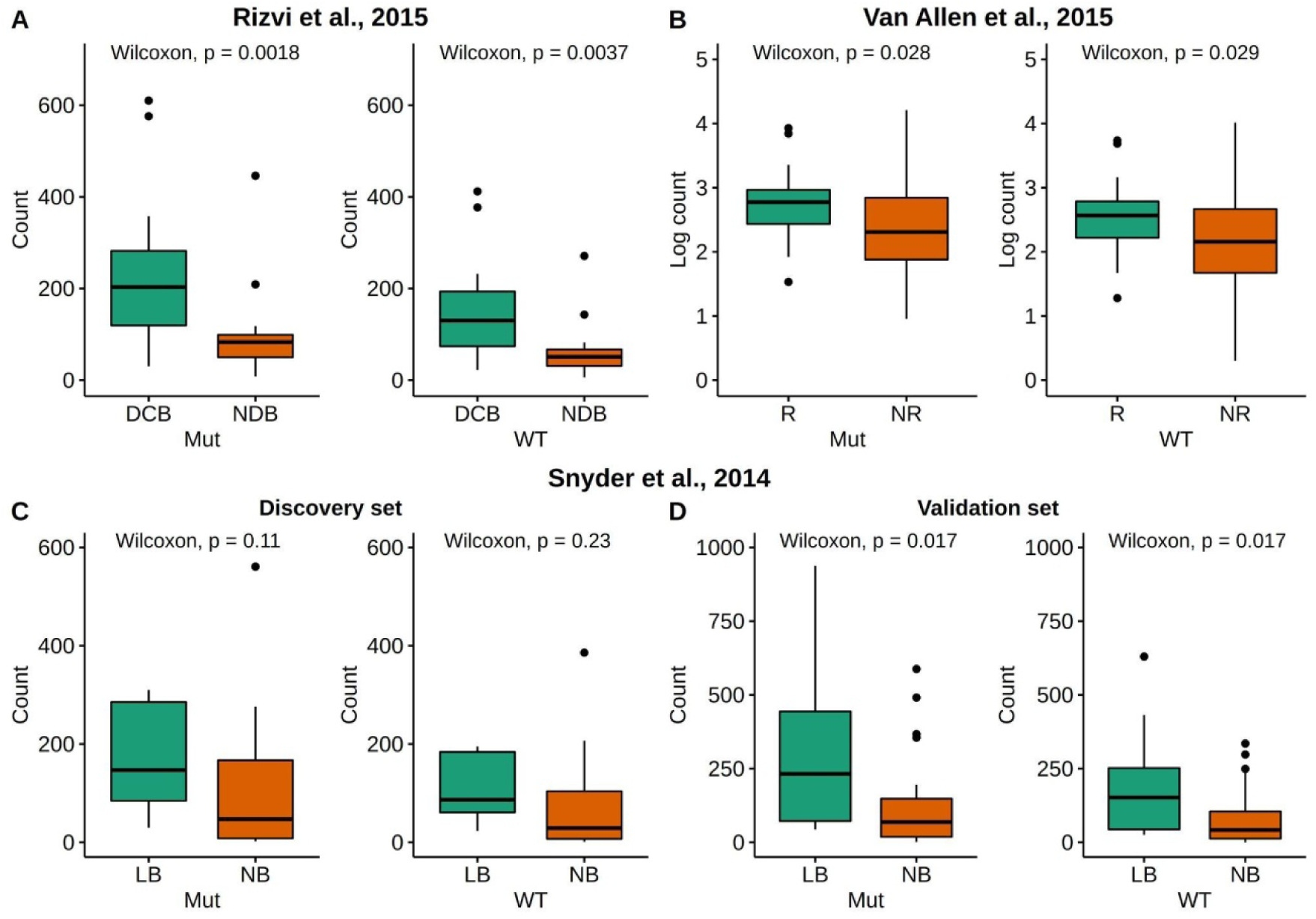
Comparison between predicted neoantigen load and clinical response for mutated and wild-type peptides. Panel **A**: Rizvi et al. clinical cohort of 34 non-small cell lung cancer patients, treated with PD-1 blockade. Panel **B**: Van Allen clinical cohort of 110 metastatic melanoma patients treated with CTLA-4 blockade. Panel **C**: Synder et al. discovery set, clinical cohort of 25 melanoma patients treated with CTLA-4 blockade. Panel **D**: Synder et al. validation set, clinical cohort of 39 melanoma patients treated with CTLA-4 blockade. DCB: Durable clinical benefit; NDB: No durable benefit; Mut: Mutated peptide; WT: Wild-type peptide; LB: Long benefit; NB: No benefit; R: Response; NR: Non-response.

Similar results were found with the Van Allen *et al*. cohort [34]. The TNB (netMHCpan version 2.4, BA<500nM) and its WT counterpart were found to be significantly associated with clinical benefit (Wilcoxon test p=0.028 and p=0.019 respectively).

Using the discovery and validation datasets from *Snyder et al*. [35] (figure 6C and 6D respectively); no significant differences were found for both TNBs (from neoantigens and WT predicted MHC-I binders - netMHCpan version 3.4) comparing responders vs. non-responders in the the discovery cohort, but when compared on the validation cohort (Wilcoxon test p=0.017) they do (predictions made with netMHCpan 4.1 see supplementary material). In addition, predicted BA of the two immunogenic neoantigens T**E**SPFEQHI and GLER**E**GFTF (validated by *Snyder et al.)* and their WT counterparts, although the first one was predicted as a SB and its WT counterpart as non-binder, the second neoantigen and its WT counterpart were both predicted as non-binders, indicating that it would not be chosen for clinical validation using the new version of netMHCpan software.

### BA scores predicted wild-type peptides as immunogenic in pre-clinical and clinical neoantigen vaccine trials

*Wolf et al*. [36] identified three top MHC-I mice neoantigens binders (FAIFNTEQ**M, V**INENYDYL, SA**Y**EKLYSL) from UVB-derived mutations on single cell melanoma mice tumors, using immunopeptidomic assays and confirmed as immunogenic by *in vivo* cytotoxicity assays. All of them were predicted here as SB (netMHCpan 4.1). The WT counterparts of the first two neoantigens were also predicted as SB and the third as WB (Supplementary Table 4).

Two validated immunogenic peptides, K**V**LKVAMKK and TA**W**QSEDSY, from a clinical trial evaluating mRNA vaccine–induced neoantigen-specific T cell immunity in gastrointestinal cancer[37], were evaluated (netMHCpan 4.1). The first, was predicted as a SB while its WT counterpart was not. But for the second, both (the mutated and its WT counterpart) were predicted as SB. Thus, the WT peptide could potentially be included in the vaccine.

## Discussion

The advent of ICB has revolutionized cancer therapy and research, succeeding in achieving durable clinical responses in patients with various cancer types such as melanoma, lung, bladder and other cancers [8].

Current scientific and clinical efforts are focused on providing both i) an accurate predictive biomarker for identifying patients who may benefit or not from ICB and ii) to identify immunogenic neoantigens that can be used in antitumor vaccination strategies. To date, there are three approved biomarkers: PD-L1 expression, microsatellite instability and/or mismatch repair deficiency, and high tumor mutation burden (H-TMB). The latter, based on the assumption that an H-TMB gives rise to neoantigens that may elicit effective anti-tumor immune response, has been approved by the FDA and widely analyzed by literature [33-35,40,41]. However, it was recently concluded that it was not a good response predictor to ICB for all cancer types [40].

Neoantigen identification, by means of the TNB, has also been evaluated as an ICB response predictor and for the development of patient-specific anticancer vaccines, leading to an unprecedented interest in the effective and accurate prediction of neoantigen immunogenicity. However, the development of efficient techniques for classification/prioritization of immunogenic neoantigens remains crucial since only a small fraction of the predicted neoantigens resulted truly immunogenic [14].

To elicit an immune response, candidate proteins holding potential neoantigens must be processed in the proteasome, then resulting neopeptides should be presented on cell surface and recognized by the TCR. Currently, most of the neoantigen immunogenicity predictors are based on two main steps after the identification of the candidate peptide: (i) prediction of binding to patient-specific HLA alleles, (ii) assessment of peptide immunogenicity based on features such as binding strength, neoantigen relative abundance and sequence similarity to viral epitopes (e.g. foreignness), among others [30,33,42].

Current immunogenic predictors were trained using the IEDB collection, which mainly contains positive-immunogenic peptides derived from non-human proteins used as positive examples. Alternatively, negative-immunogenic peptides were generated by random sampling of peptides from the human proteome or by building random amino acid sequences. Therefore, they were mainly developed to evaluate foreign peptides instead of self-derived ones such as neoantigens. The limitations of the current concepts and methods, based on BA and/or derived scores, to classify neoantigens/WT peptides as immunogenic or non-immunogenic were exposed, failing to accurately classify/prioritize immunogenic neoantigens. In addition, it was found that WT counterparts peptides were also classified as immunogenic, which is in agreement with previously reported evidence that natural derived peptides can be presented by the MHC-I molecule and trigger an immune reaction [3,16], spotlighting the use of natural derived peptides as negative immunogenic examples is a misleading strategy if they have not been experimentally confirmed as true negatives. Similarly, this may also explain the prediction inaccuracy of current approaches for the recognition of true negative immunogenic neoantigens, consequently resulting in high false positive rates.

This observation reinforces the utility of a neoantigen curation effort to provide a new database holding, to date, 61 nine-mer neoantigens with experimentally proven binding to the MHC-I molecule, as well as their positive or negative immunogenicity. In this way it provides a truly immunogenic and non-immunogenic neoantigen database. It is expected that current and future versions of ITSNdb may serve as a benchmark dataset to test neoantigen immunogenicity predictors. It will allows the recognition of truly negative neoantigens, facilitating the development of more accurate bioinformatic tools and their performance evaluation and prediction confidence. In this regards, the use of the DOP seems to be better than the Youden method to identify classification thresholds. It optimizes sensitivity and confidence of both positive and negative predictions.

The use of the ITSNdb revealed that the high false positive rate of current methods limits their application in the selection or prioritization of patient-specific neoantigens for vaccination strategies. Furthermore, the identification of WT peptides as candidate neoantigens, may have a negative impact on current preclinical and clinical trials of antitumoral mRNA neoantigens-derived vaccines [36,37].

Our results on immunogenic prediction for both mutated and WT counterpart pairs substantiates the hypothesis that natural human protein fragments may have immunogenic potential that are exacerbated by just one amino acid substitution. Further research in this area is warranted.

Another important finding was that the TNB, established upon current neoantigen definition (based on BA to the MHC-I), failed to overcome the TMB as ICB response predictor. Additionally, TNB estimated over predictions from neoantigens WT counterparts, was also associated with the ICB response. These findings on WT peptides, suggest that the concept of “foreignness” is not a predictor of immunogenicity, in agreement with previous reports concluding that similarity, or not, to viral peptides was poorly informative to predict/prioritize neoantigen immunogenicity [30].

## Conclusions

The ITSNdb is a very promising database to evaluate current and future tools for the prediction of neoantigen immunogenicity. Its benchmark use indicates that new rules and novel measures/metrics should be devised to improve the performance on predicting or prioritizing immunogenic neoantigens in order to become a valid screening tool for both accurate predictive biomarker of ICB responses as well as in anticancer vaccine development. It is expected that current and future versions of this novel curated neoantigen database may constitute a valid benchmark for upcoming neoantigens and immunogenicity identification together with the proposed DOP metric for model selection/evaluation.

### Key points

- A new neoantigen database, ITSNdb, with true immunogenic and non-immunogenic neoantigens, is presented to provide a new tool for fair prediction software evaluation
- A comparative benchmark between the state of the art tools to predict immunogenic neoantigens, using ITSNdb, reveals that new rules should be discovered and devised
- The clinical impact on tumor neoantigen burden of the current immunogenic neoantigen prediction was evaluated on ICB Clinical trials
- Current methods predict a high proportion of wild type neoantigens counterpart peptides as immunogenic with impact in neoantigens vaccine and biomarkers design

## Data availability

The data underlying this article are available in the article and in its online supplementary material.

## Acknowledgments

We want to thank Dr. Ignacio Wichman for his help reading and discussing this work; and Dr. Raphael Trevizani for providing us deepitope and explaining to us how to run it.

## Supplementary data

Supplementary material

Supplementary table 1

Supplementary table 2

Supplementary table 3

Supplementary table 4

