## supplementary material for "MHC-I binding affinity derived metrics fail to predict tumor specific neoantigen immunogenicity"

1- Centro de Investigación y Desarrollo en Inmunología y Enfermedades Infecciosas (CIDIE), Consejo Nacional de Investigaciones Científicas y Técnicas (CONICET)/Universidad Católica de Córdoba (UCC). X5016HDK. Córdoba, Argentina.

2- Laboratorio de Inmuno Oncologia Traslacional (Instituto de Biología y Medicina Experimental - IBYME) – CONICET.

3- Instituto Académico Pedagógico de Ciencias Básicas y Aplicadas, Universidad Nacional de Villa María, Villa María, Córdoba X5900, Argentina.

4- Departamento de Bioquímica Clínica, Facultad de Ciencias Químicas, Universidad Nacional de Córdoba, Córdoba, Argentina.

5- Consejo Nacional de Investigaciones Científicas y Técnicas, Centro de Investigaciones en Bioquímica Clínica e Inmunología, Córdoba, Argentina.

6- Facultad de Ciencias de la Salud, Universidad Católica de Córdoba (UCC). X5016HDK. Córdoba, Argentina.

7- Facultad de Ingeniería. Universidad Católica de Córdoba. X5016HDK. Córdoba, Argentina.

8- Facultad de Ingeniería, FCEFyN Universidad Nacional de Córdoba (UNC). X5016. Córdoba, Argentina.

### Detailed methods

#### Data collection

The neoantigens were first collected from PubMedTM using “neoantigen" or “neoepitopes” as keywords. The resulting publications were manually curated. Those neoantigens whose inclusion criteria were explicitly described were included in the new database. Next, references from the primary publications (i.e., those that were found using the keywords) were carefully revised. If additional peptides that met the defined criteria were found in these references, the curation process was repeated. The references of the neoantigens from DbPepNeo and CAPD were also examined and curated. At the end of this process, neoantigens from 19 scientific publications were included from more than 50 revised manuscripts.

#### Data exclusion

Scientific publications were curated looking for neoantigens and their immunogenicity. It should have come from SNVs, and mutated and wild type peptides sequences of 9 AA were collected.

When analyzing the possible incorporation of one peptide into ITSNdb, first there had to be an explicit description of positive experimental binding assays, otherwise it was discharged. In second place, immunogenic assays should be performed, bringing either positive or negative results; if there was no experimental validation of immunogenicity, the peptide was not included into the database. Finally all wild type sequences were searched into the origin protein sequence (referenced into the paper). Five peptides out of 66 were excluded from the database; Two of them did not appear into the protein sequence: CLSPQTLAA (PRKCDBP), YISKCWDHA (HSD17B7); One had an origin gene that was named provisionally as MUM3 and there is no record about what is it actual gene: EASIQPITR; and the last two, their sequence were changed when synthesized in order to improve the binding for a vaccine: AMFRSVPTV (TKT), HPYASLSRV (MRPS5).

#### Software and algorithms application

Seven predictive software were run over the 61 peptides from ITSNdb, in order to compare software predictions with validated data. Statistical analysis was performed with R language version 3.6.3.

#### Key resources table

| Software | URL | Reference |
|---|---|---|
| R 3.6.3 | CRAN | https://www.r-project.org/ |
| NetMHCpan 4.1 | https://services.healthtech.dtu.dk/service.php?NetMHCpan-4.1 | https://academic.oup.com/nar/article/48/W1/W449/5837056 |
| MHCflurry 2.0 | https://github.com/openvax/mhcflurry | https://www.sciencedirect.com/science/article/pii/S2405471220302398 |
| MixMHCpred 2.1 | https://github.com/GfellerLab/MixMHCpred | https://www.jimmunol.org/content/201/12/3705 |
| Deepimuno | https://deepimmuno.research.cchmc.org/ | https://academic.oup.com/bib/article/22/6/bbab160/6261914 |
| CIImm | http://tools.iedb.org/immunogenicity/ | https://journals.plos.org/ploscompbiol/article?id=10.1371/journal.pcbi.1003266 |
| Deepitope | | |
| PRIME | http://prime.gfellerlab.org/ | https://www.sciencedirect.com/science/article/pii/S2666379121000057 |

#### Software description

##### BA prediction software

NetMHCpan 4.1: is an artificial neural network, designed to predict peptide-MHC-I binding. It uses the NNAlign_MA framework to allow for the incorporation of eluted mass spectrometry (MS) data. Thus, it is trained to estimate both the BA and the eluted mass as ranks. For a peptide-HLA restriction pair, it provides the eluted rank (%R) and the BA (nM) to their restricted HLA [22]. The authors propose to classify the peptides as either strong (SB), weak (WB) or non-binders, according to predefined score thresholds (by default, SB=%R ≤ 0.5% and WB=0.5%<%R≤ 2%).

MHCflurry 2.0: is a trained convolutional neural network (CNN) that includes affinity measurements and MS datasets. It predicts BA in nM (lower values indicate stronger affinity and the percentile (P) of the affinity prediction (calculated among a large number of random peptides tested on that allele). The authors do not provide predefined thresholds to outline strong binders, but they suggest using the popular threshold of BA <500 nM or P<2% to differentiate binder from non-binder peptides [23].

MixMHCpred 2.1: is a binding motif deconvolution in HLA-I peptidomic model, trained on mass spectrometry elution data. It provides affinity scores (the higher the score, the stronger the peptide binding to the HLA-I) and ranks (the lower the rank, the stronger the peptide binding to the HLA-I) [24].

#### Immunogenicity prediction software

DeepImmuno-CNN 1.2: A web application implementing a CNN that predicts immunogenicity of MHC–peptide pairs. It uses MHCflurry to predict BA. The model provides a continuous immunogenic score. The greater the peptide scores, the higher the chance for it to be immunogenic. No score threshold is provided to define immunogenicity [25].

T cell class I pMHC immunogenicity predictor (CIImm): A web application implementing a model to predict the immunogenicity of new peptides-MHCs complexes. It provides an immunogenicity score calculated as a position-dependent weighted sum of the non-anchor amino acids of the neoantigen amino acid sequence. Higher scores mean a higher chance for the peptide to be immunogenic. No threshold is provided to define peptide immunogenicity [26].

Deepitope: is a CNN trained with IEDB data to classify peptides as immunogenic or non-immunogenic.

PRIME 1.0: web application implementing a class I immunogenicity predictor combining affinity predictions to HLA-I molecules, performed by mixMHCpred, together with TCR recognition propensity, trained as a logistic regression. Non-immunogenic peptides receive a score of 0 [27].

#### Utilized code and web applications

##### netMHCpan:

Run from R

```
> netMHC <- system2("/path/to/netMHC/netMHCpan-4.1/netMHCpan", args = c("-p", "-a", allele, pep.temp, "-BA"), stdout = TRUE)
```

##### mixMHCpred:

Run from R

```
> system2("/path/to/MixMHCpred-2.1/MixMHCpred", args = c("-i", pep.temp, "-o", "mixout.txt", "-a", allele), stdout = TRUE)
```

##### MHCflurry:

```
$ mhcflurry-predict INPUT.csv –out RESULT.csv
```

##### Deepitope

```
$ python src/example.py   model/   peptides1.txt   peptides2.txt
```

##### CIImm:

Web application: http://tools.iedb.org/immunogenicity/ (run with default options)

##### Deepimmune:

Web application: https://deepimmuno.research.cchmc.org/ (run with default options)

##### PRIME:

Web application: http://prime.gfellerlab.org/ (run with default options)

#### Immunotherapy datasets

Publicly available datasets from three different publications were collected. ICB response over 34 non–small cell lung cancer patients treated with pembrolizumab, an antibody targeting programmed cell death protein-1 (PD-1) was analyzed by *Rizvi et al* [33]; a cohort of 110 patients with metastatic melanoma treated with CTLA-4 blockade and their treatment response association, analyzed by Van Allen *et al*. [34]; and another two subject cohorts with metastatic melanoma treated with anti-CTA4 ICB therapy and their response over 25 and 39 patients, evaluated by Snyder *et al*. [35]. All datasets contain mutated peptide and WT counterpart sequences and BA data.

### Supplementary figure 1

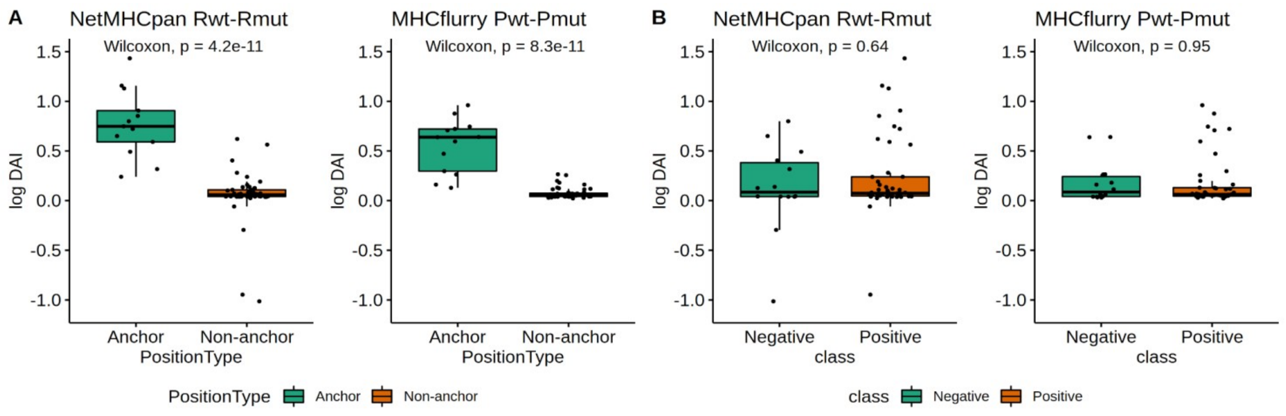


***Fig S1.*** *Comparison between DAI values for netMHCpan and MHCflurry wild type and mutated peptides rank/percentile differenceA) DAI differences vs. position type (anchor vs non-anchor) and B) DAI difference vs. immunological class (positive vs negative). DAI were calculated as (logDAI) for visualization purposes.*

### Supplementary figure 2

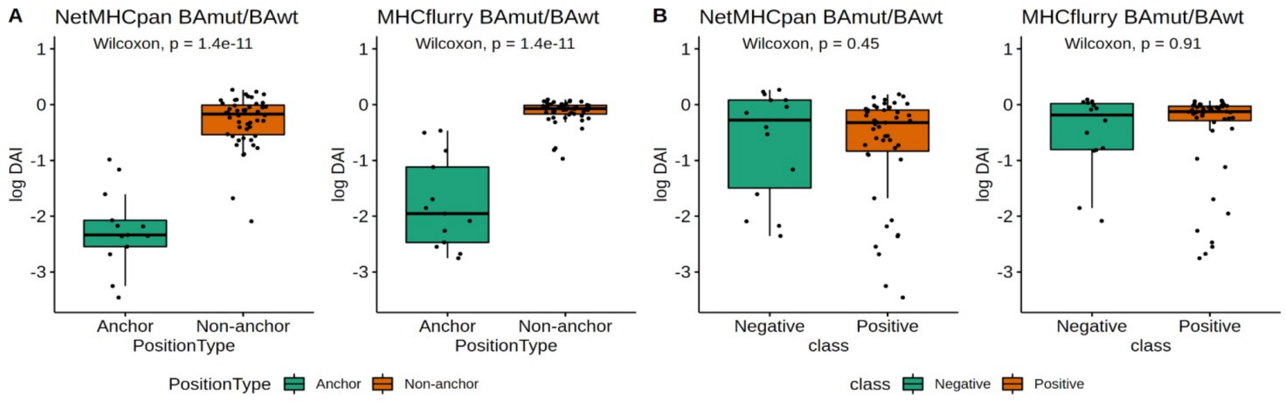


***Fig S2****. Comparison between DAI values for netMHCpan and MHCflurry wild type and mutated peptides binding affinity ratio. A) DAI differences vs. position type (anchor vs non-anchor) and B) DAI difference vs. immunological class (positive vs negative). DAI were calculated as (logDAI) for visualization purposes.*

### Supplementary figure 3

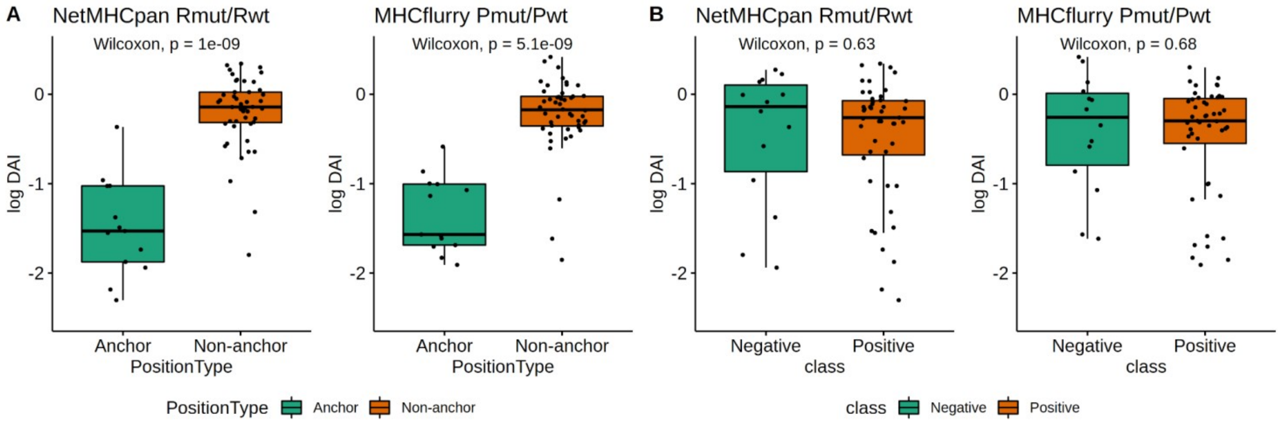


***Fig S3.*** *Comparison between DAI values for netMHCpan and MHCflurry wild type and mutated peptides rank/percentile ratio, A) DAI differences vs. position type (anchor vs non-anchor) and B) DAI difference vs. immunological class (positive vs negative). DAI were calculated as (logDAI) for visualization purposes.*

### Supplementary figure 4

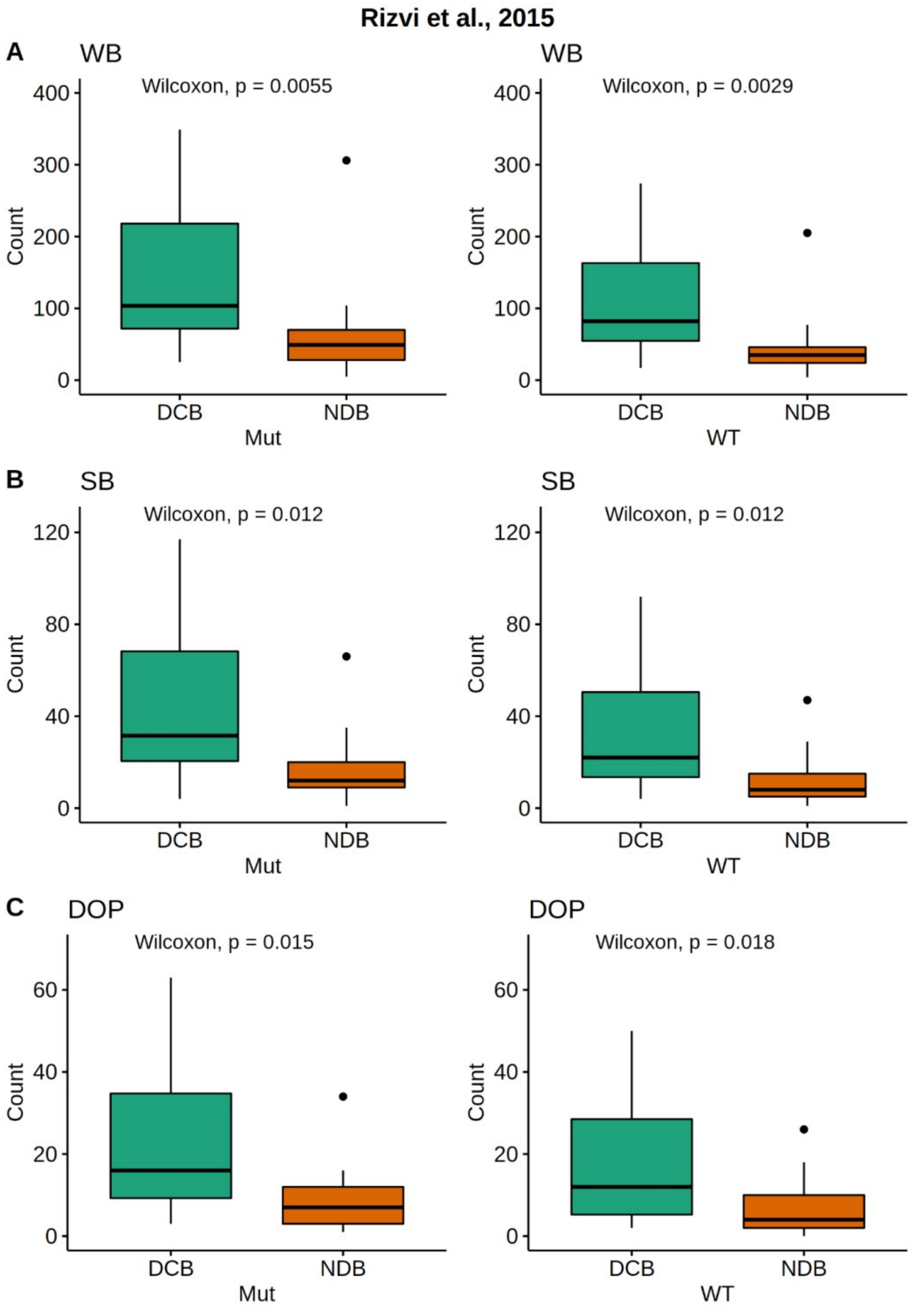


***Fig S4.*** *Comparison between the amount of netMHCpan 4.1 predicted neoantigen burden for ICB durable clinical benefit (DCB) vs. No durable clinical benefit (NDB) patients from the Rizvi et al. dataset. Left panels represent netMHCpan 4.1 neoantigen prediction on mutated papetides (TNB) and right panels represent neoantigen burden calculated over netMHCpan 4.1 wild type counterpart predictions. A) using weak binders threshold B) using strong binders threshold and c) using DOP threshold.*

### Supplementary figure 5

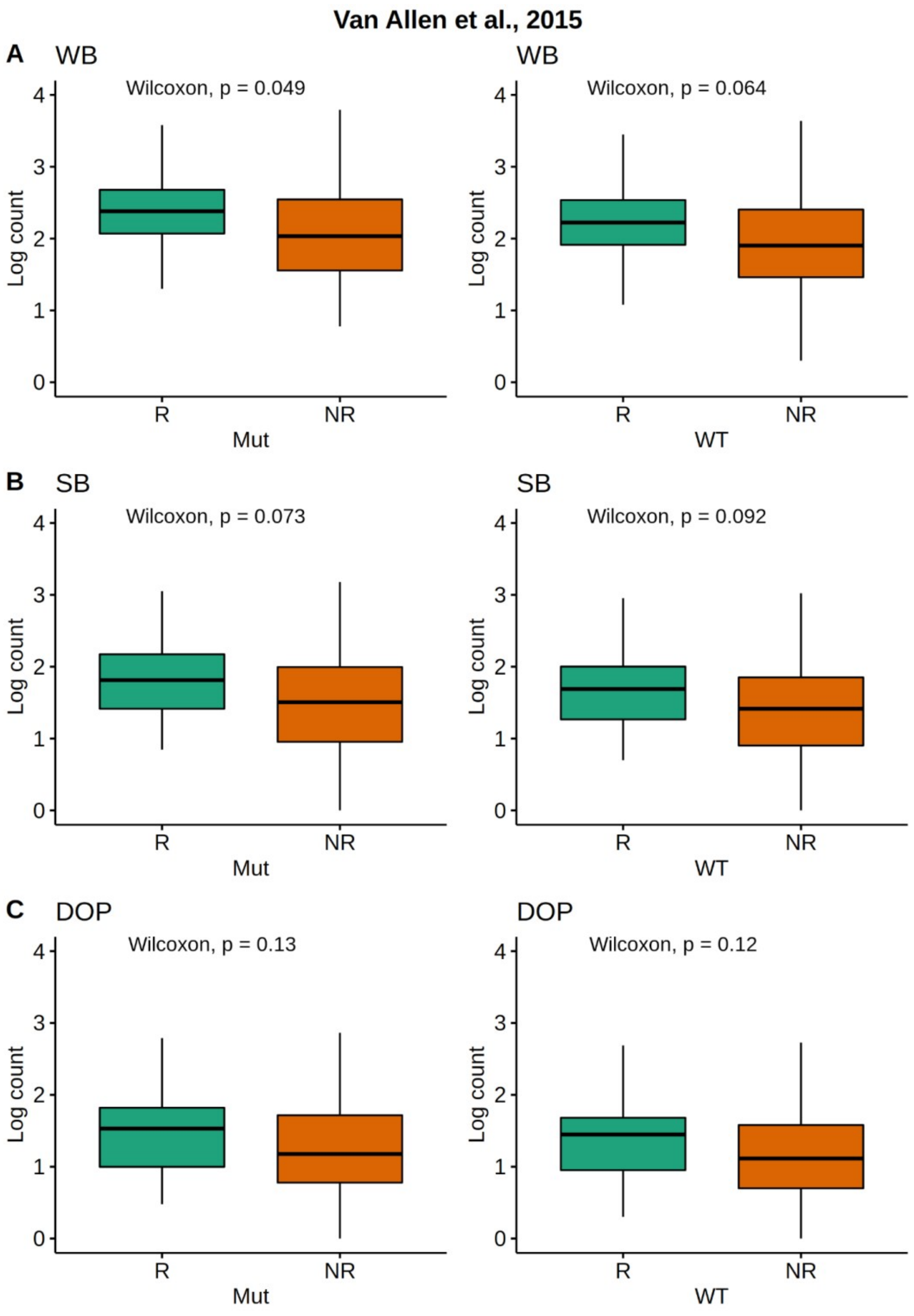


***Fig S5.*** *Comparison between the amount of netMHCpan 4.1 predicted neoantigen burden for ICB responders (R) vs. non responders (NR) patients from the Van Allen et al. dataset. Left panels represent netMHCpan 4.1 neoantigen prediction on mutated papetides (TNB) and right panels represent neoantigen burden calculated over netMHCpan 4.1 wild type counterpart predictions. A) using weak binders threshold B) using strong binders threshold and c) using DOP threshold.*

### Supplementary figure 6

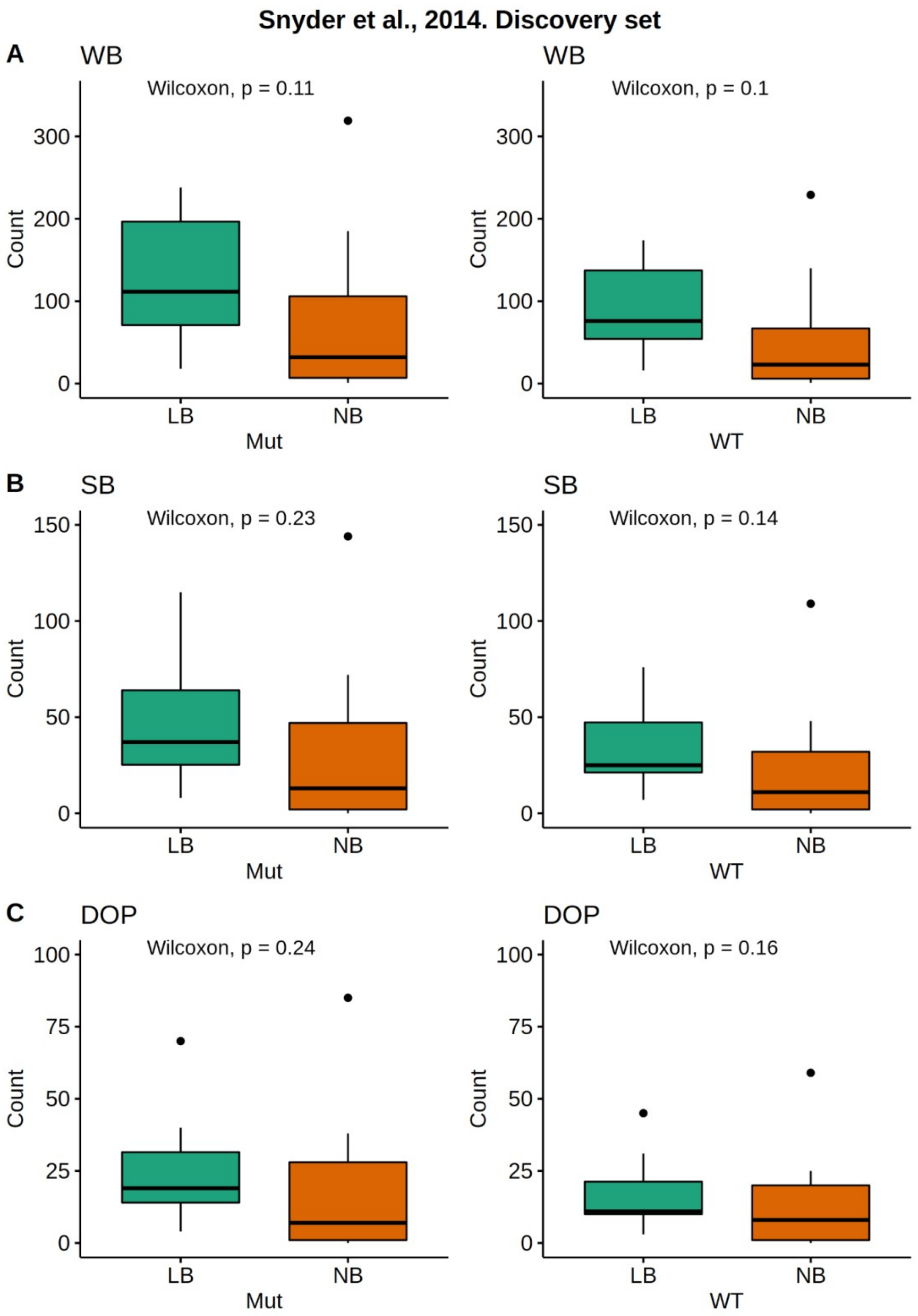


***Fig S6.*** *Comparison between the amount of netMHCpan 4.1 predicted neoantigen for Long Benefit (LB) vs. No Benefit (NB) ICB treatment according to clinical response for mutated peptides (Mut) or TNB and wild type (WT) counterparts from the Snyder et al. discovery dataset. A) using weak binders threshold B) using strong binders threshold and c) using DOP threshold.*

### Supplementary figure 7

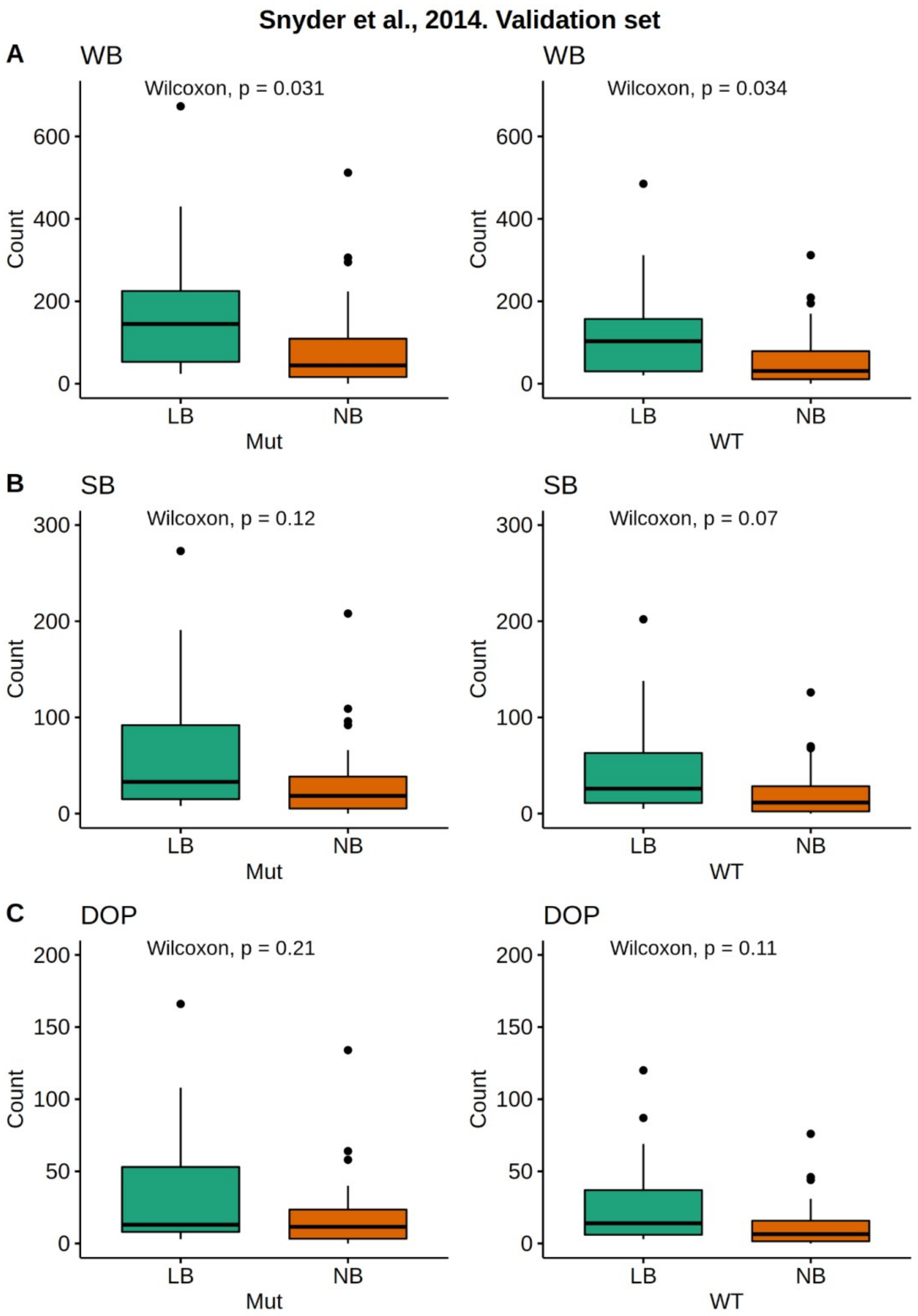


***Fig S7.*** *Comparison between the amount of netMHCpan 4.1 predicted neoantigen for Long Benefit (LB) vs. No Benefit (NB) ICB treatment clinical response for mutated peptides (Mut) or TNB and wild type (WT) counterparts from the Snyder et al. validation dataset. A) using weak binders threshold B)using strong binders threshold and c) using DOP threshold.*
