## Supplementary table 1 for "MHC-I binding affinity derived metrics fail to predict tumor specific neoantigen immunogenicity"

**ITSNdb**

| **Paper** | **Author** |
|---|---|
| https://www.jimmunol.org/content/160/12/6188.long | Gueguen;Fritsch |
| https://www.ncbi.nlm.nih.gov/pmc/articles/PMC4607110/?report=reader#!po=65.3846 | Cohen |
| https://science.sciencemag.org/content/352/6291/1337 | Stronen |
| https://www.ncbi.nlm.nih.gov/pmc/articles/PMC4607110/?report=reader#!po=65.3846 | Cohen |
| https://science.sciencemag.org/content/352/6291/1337 | Stronen |
| https://www.ncbi.nlm.nih.gov/pmc/articles/PMC4549796/ | Carreno |
| https://www.ncbi.nlm.nih.gov/pmc/articles/PMC3757932/#SD2 | Robbins;Fritsch |
| https://www.ncbi.nlm.nih.gov/pmc/articles/PMC4549796/ | Carreno |
| https://www.ncbi.nlm.nih.gov/pmc/articles/PMC2193899/ | Akatsuka;Fritsch |
| https://www.nature.com/articles/ncomms13404#MOESM1316 | Bassani-Sternberg |
| https://cancerres.aacrjournals.org/content/58/22/5144.long | Hogan;Fritsch |
| https://cancerres.aacrjournals.org/content/61/9/3718.long | Karanikas;Fritsch |
| https://www.ncbi.nlm.nih.gov/pmc/articles/PMC4607110/?report=reader#!po=65.3846 | Cohen |
| https://www.ncbi.nlm.nih.gov/pmc/articles/PMC4549796/ | Carreno |
| https://www.ncbi.nlm.nih.gov/pmc/articles/PMC4607110/?report=reader#!po=65.3846 | Cohen |
| https://cancerres.aacrjournals.org/content/59/22/5785.full | Chiari |
| https://www.ncbi.nlm.nih.gov/pmc/articles/PMC4549796/ | Carreno |
| https://www.ncbi.nlm.nih.gov/pmc/articles/PMC2175172/#!po=50.0000 | Zhou |
| https://www.ncbi.nlm.nih.gov/pmc/articles/PMC3757932/#SD2 | Robbins |
| https://www.ncbi.nlm.nih.gov/pmc/articles/PMC6453117/pdf/nihms-999473.pdf | Parkhurst |
| https://www.ncbi.nlm.nih.gov/pmc/articles/PMC6453117/pdf/nihms-999473.pdf | Parkhurst |
| https://www.nature.com/articles/ncomms13404#MOESM1316 | Bassani-Sternberg |
| https://www.ncbi.nlm.nih.gov/pmc/articles/PMC4549796/ | Carreno |
| https://science.sciencemag.org/content/352/6291/1337 | Stronen |
| https://www.ncbi.nlm.nih.gov/pmc/articles/PMC3757932/#SD2 | Robbins;Fritsch |
| https://www.ncbi.nlm.nih.gov/pmc/articles/PMC4607110/?report=reader#!po=65.3846 | Cohen |
| https://www.ncbi.nlm.nih.gov/pmc/articles/PMC4549796/ | Carreno |
| https://www.ncbi.nlm.nih.gov/pmc/articles/PMC1266037/ | Lennerz;Fritsch |
| https://www.ncbi.nlm.nih.gov/pmc/articles/PMC4549796/ | Carreno |
| https://www.nature.com/articles/ncomms13404#MOESM1316 | Bassani-Sternberg |
| https://www.nature.com/articles/ncomms13404#MOESM1316 | Bassani-Sternberg |
| https://www.ncbi.nlm.nih.gov/pmc/articles/PMC4549796/ | Carreno |
| https://science.sciencemag.org/content/352/6291/1337 | Stronen |
| https://www.ncbi.nlm.nih.gov/pmc/articles/PMC4549796/ | Carreno |
| https://www.ncbi.nlm.nih.gov/pmc/articles/PMC3757932/#SD2 | Robbins;Fritsch |
| https://www.ncbi.nlm.nih.gov/pmc/articles/PMC2175172/#!po=50.0000 | Zhou |
| https://science.sciencemag.org/content/352/6291/1337 | Stronen |
| https://www.nature.com/articles/s41467-018-06405-9 | Zhang |
| https://science.sciencemag.org/content/352/6291/1337 | Stronen |
| https://www.ncbi.nlm.nih.gov/pmc/articles/PMC4549796/ | Carreno |
| https://www.ncbi.nlm.nih.gov/pmc/articles/PMC4549796/ | Carreno |
| https://www.nature.com/articles/ncomms13404#MOESM1316 | Bassani-Sternberg |
| https://www.ncbi.nlm.nih.gov/pmc/articles/PMC4549796/ | Carreno |
| https://science.sciencemag.org/content/352/6291/1337 | Stronen |
| https://www.ncbi.nlm.nih.gov/pmc/articles/PMC4549796/ | Carreno |
| https://www.ncbi.nlm.nih.gov/pmc/articles/PMC4549796/ | Carreno |
| https://science.sciencemag.org/content/352/6291/1337 | Stronen |
| https://cancerres.aacrjournals.org/content/65/3/1079.long | Zhou;Fritsch |
| https://www.ncbi.nlm.nih.gov/pmc/articles/PMC4549796/ | Carreno |
| https://www.ncbi.nlm.nih.gov/pmc/articles/PMC1266037/ | Lennerz;Fritsch |
| https://www.ncbi.nlm.nih.gov/pmc/articles/PMC6453117/pdf/nihms-999473.pdf | Parkhurst |
| https://www.ncbi.nlm.nih.gov/pmc/articles/PMC1266037/ | Lennerz |

| | |
|---|---|
| https://www.nature.com/articles/ncomms13404#MOESM1316 | Bassani-Sternberg |
| https://www.ncbi.nlm.nih.gov/pmc/articles/PMC6453117/pdf/nihms-999473.pdf | Parkhurst |
| https://www.ncbi.nlm.nih.gov/pmc/articles/PMC4607110/?report=reader#!po=65.3846 | Cohen;Parkhurst |
| https://pubmed.ncbi.nlm.nih.gov/24891321/ | Rajasagi |
| https://pubmed.ncbi.nlm.nih.gov/9461441/ | den Haan;fritsch |
| https://www.ncbi.nlm.nih.gov/pmc/articles/PMC4549796/ | Carreno |
| https://pubmed.ncbi.nlm.nih.gov/24891321/ | Rajasagi;Fritsch |
| https://www.nature.com/articles/ncomms13404#MOESM1316 | Bassani-Sternberg |
| https://www.ncbi.nlm.nih.gov/pmc/articles/PMC3757932/#SD2 | Robbins;Fritsch |

| Sequence | WT | mutPosition | PositionType | Gene | Tumor | HLA | class |
|---|---|---|---|---|---|---|---|
| AEPINIQTW | AEPIDIQTW | 5 | Non-anchor | KIAA0205 | Bladder | HLA-B44:03 | Positive |
| ALIHHNTYL | ALIHHNTHL | 8 | Non-anchor | ERBB2 | Melanoma | HLA-A02:01 | Positive |
| ALSPVIPHI | ALSPVIPLI | 8 | Non-anchor | MLL2 | Melanoma | HLA-A02:01 | Positive |
| ALYGFVPVL | ALYGSVPVL | 5 | Non-anchor | GANAB | Melanoma | HLA-A02:01 | Positive |
| AVGSYVYSV | AVGSHVYSV | 5 | Non-anchor | PGM5 | Melanoma | HLA-A02:01 | Positive |
| AVIDAYTEI | AVINAYTEI | 4 | Non-anchor | LARP7 | Melanoma | HLA-A02:01 | Negative |
| CILGKLFTK | CILGELFTK | 5 | Non-anchor | CDK12 | Melanoma | HLA-A11:01 | Positive |
| CLNEYHLFL | CLNEYHLFF | 9 | Anchor | TMEM48 | Melanoma | HLA-A02:01 | Positive |
| DYLQYVLQI | DYLQCVLQI | 5 | Non-anchor | BCL2A1 | AML | HLA-A24:02 | Positive |
| ETSKQVTRW | ETSEQVTRW | 4 | Non-anchor | GABPA | Melanoma | HLA-A25:01 | Negative |
| ETVSEQNSV | ETVSEESNV | 8 | Non-anchor | EF2 | Lung | HLA-A68:01 | Positive |
| FLDEFMEGV | FLDEFMEAV | 8 | Non-anchor | ME-1 | Lung | HLA-A02:01 | Positive |
| FLIYLDVSV | FLTYLDVSV | 3 | Non-anchor | WDR46 | Melanoma | HLA-A02:01 | Positive |
| FLYNLLTRV | FLYNPLTRV | 5 | Non-anchor | SEC24A | Melanoma | HLA-A02:01 | Positive |
| FMPDFDLHL | SMPDFDLHL | 1 | Non-anchor | AHNAK | Melanoma | HLA-A02:01 | Positive |
| FRSGLDSYV | FRSRLDSYV | 4 | Non-anchor | MUM-2 | Melanoma | HLA-C06:02 | Positive |
| FVSALCMFL | FVSALRMFL | 6 | Non-anchor | ARFGEF1 | Melanoma | HLA-A02:01 | Negative |
| GIVEGLITT | GIVEGLMTT | 7 | Non-anchor | GAPDH | Melanoma | HLA-A02:01 | Positive |
| GLFGDIYLA | GSFGDIYLA | 2 | Anchor | CSNK1A1 | Melanoma | HLA-A02:01 | Positive |
| GLLDEDFYA | GPLDEDFYA | 2 | Anchor | UGGT2 | Melanoma | HLA-A02:01 | Positive |
| GQFLTPNSH | GQFLAPNSH | 5 | Non-anchor | TFDP2 | Melanoma | HLA-B15:01 | Positive |
| GRIAFFLKY | GRIAFSLKY | 6 | Non-anchor | SYTL4 | Melanoma | HLA-B27:05 | Positive |
| IILVAVPHV | IILVAVQHV | 7 | Non-anchor | EXOC8 | Melanoma | HLA-A02:01 | Positive |
| ILAALFLGL | IWAALFLGL | 2 | Anchor | SLC38A1 | Melanoma | HLA-A02:01 | Positive |
| ILNAMIAKI | ILNAMITKI | 7 | Non-anchor | HAUS3 | Melanoma | HLA-A02:01 | Positive |
| ILTGLNYEV | ILTGLNYEA | 9 | Anchor | NSDHL | Melanoma | HLA-A02:01 | Positive |
| IMAHCILDL | IIAHCILDL | 2 | Anchor | SRP9 | Melanoma | HLA-A02:01 | Negative |
| KIFSEVTLK | KIFSEVTPK | 8 | Non-anchor | SIRT2 | Melanoma | HLA-A03:01 | Positive |
| KLANPLPYT | KLAKPLPYT | 4 | Non-anchor | SMOX | Melanoma | HLA-A02:01 | Negative |
| KLILWRGLK | KPILWRGLK | 2 | Anchor | NCAPG2 | Melanoma | HLA-A03:01 | Positive |
| KLKLPIIMK | KLKLPMIMK | 6 | Non-anchor | AKAP6 | Melanoma | HLA-A03:01 | Negative |
| KLMNIQQKL | KLMNIQQQL | 8 | Non-anchor | AKAP13 | Melanoma | HLA-A02:01 | Positive |
| KLSHQLVLL | KLSHQPVLL | 6 | Non-anchor | SNX24 | Melanoma | HLA-A02:01 | Positive |
| KMIGNHLWV | EMIGNHLWV | 1 | Non-anchor | CDKN2A | Melanoma | HLA-A02:01 | Positive |
| KTLTSVFQK | ETLTSVFQK | 1 | Non-anchor | MATN2 | Melanoma | HLA-A11:01 | Positive |
| LADEAEVYL | LADEAEVHL | 8 | Non-anchor | GAS7 | Melanoma | HLA-A02:01 | Positive |
| LIIPFIHLI | LIIPCIHLI | 5 | Non-anchor | USP28 | Melanoma | HLA-A02:01 | Positive |
| LLDGFLATV | LLDRFLATV | 4 | Non-anchor | CCNI | Melanoma | HLA-A02:01 | Positive |
| LLFGMPPCL | LLFGMTPCL | 6 | Non-anchor | MRM1 | Melanoma | HLA-A02:01 | Positive |
| LLSIIFFPA | LLSIISFPA | 6 | Non-anchor | PHKA2 | Melanoma | HLA-A02:01 | Negative |
| LLSIVPCTV | LLSIVLCTV | 6 | Non-anchor | GPX8 | Melanoma | HLA-A02:01 | Negative |
| LPIQYEPVL | PPIQYEPVL | 1 | Non-anchor | SEC23A | Melanoma | HLA-B35:03 | Negative |
| MLGEQLFPL | MLGERLFPL | 5 | Non-anchor | PABPC1 | Melanoma | HLA-A02:01 | Positive |
| NLNCCSVPV | NLNRCSVPV | 4 | Non-anchor | GNL3L | Melanoma | HLA-A02:01 | Positive |
| QLDKCSAFV | QLDQCSAFV | 4 | Non-anchor | UTRN | Melanoma | HLA-A02:01 | Negative |
| QLSCISTYV | QLSCTSTYV | 5 | Non-anchor | OR8B3 | Melanoma | HLA-A02:01 | Positive |
| QQFAVGSYV | QQFAVGSHV | 8 | Non-anchor | PGM5 | Melanoma | HLA-A02:01 | Positive |
| QTACEVLDY | QTTCEVLDY | 3 | Non-anchor | KIAA1440 | Renal | HLA-A01:01 | Positive |
| QTIDNIVFL | QTIDNIVFF | 9 | Anchor | ARFGEF1 | Melanoma | HLA-A02:01 | Negative |
| RPHVPESAF | GPHVPESAF | 1 | Non-anchor | RBAF | Melanoma | HLA-B07:02 | Positive |
| RVSTLRVSL | RVSTLRVSP | 9 | Anchor | GNB5 | Melanoma | HLA-B07:02 | Positive |
| SHETVIIEL | SHETVTIEL | 6 | Non-anchor | SNRPD1 | Melanoma | HLA-B38:01 | Positive |

| | | | | | | | |
|---|---|---|---|---|---|---|---|
| SPGPVKLEL | SPGPVKLEP | 9 | Anchor | NOP16 | Melanoma | HLA-B07:02 | Negative |
| SYMIMEIEL | SSMIMEIEL | 2 | Anchor | FBXO21 | Melanoma | HLA-C14:02 | Positive |
| TLWCSPIKV | TPWCSPIKV | 2 | Anchor | SRPX | Melanoma | HLA-A02:01 | Positive |
| TPTVPSSSF | TPTVPSGSF | 7 | Non-anchor | ALMS1 | CLL | HLA-B35:01 | Positive |
| VLHDDLLEA | VLRDDLLEA | 3 | Non-anchor | KIAA0223 | AML | HLA-A02:01 | Positive |
| VLLRALPVL | VLLRALPVP | 9 | Anchor | MRPS17 | Melanoma | HLA-A02:01 | Negative |
| VVMSWAPPV | VVLSWAPPV | 3 | Non-anchor | FNDC3B | CLL | HLA-A02:01 | Positive |
| YIDERFERY | YIDEQFERY | 5 | Non-anchor | SEPT2 | Melanoma | HLA-A01:01 | Negative |
| YTDFHCQYV | YTDFPCQYV | 5 | Non-anchor | PPP1R3B | Melanoma | HLA-A01:01 | Positive |

| netMHCpan_BAmut | netMHCpan_BAWT | netMHCpan_Rmut | netMHCpan_RWT | MHCflurry_BAmut |
|---|---|---|---|---|
| 83.57 | 107.36 | 0.0188 | 0.0178 | 56.5360112056013 |
| 17.83 | 95.27 | 0.0579 | 0.1235 | 14.8245941743442 |
| 11.47 | 11.07 | 0.0077 | 0.0158 | 11.6796746607546 |
| 8.06 | 15.72 | 0.0216 | 0.0257 | 15.007109781573 |
| 14.07 | 38.55 | 0.1353 | 0.0962 | 15.1525507243698 |
| 270.97 | 920.02 | 0.514 | 1.954 | 44.4606139579126 |
| 21.55 | 28.5 | 0.2786 | 0.3867 | 29.5713121623323 |
| 17.08 | 2020.25 | 0.4711 | 4.9778 | 31.3390998222739 |
| 174.1 | 239.52 | 0.1651 | 0.3511 | 57.9581377880844 |
| 518.29 | 281.07 | 0.0094 | 0.005 | 34.8964567219062 |
| 5714.73 | 8911.96 | 5.7873 | 8.3556 | 3824.38870331386 |
| 2.95 | 2.74 | 0.021 | 0.0105 | 13.4708834234424 |
| 3.53 | 3.94 | 0.1231 | 0.15 | 13.9859616803895 |
| 3.05 | 3.82 | 0.011 | 0.005 | 10.5595226556969 |
| 6.98 | 36.95 | 0.0568 | 0.1293 | 14.3119061414911 |
| 46.49 | 57.97 | 0.0999 | 0.1515 | 68.0631092383503 |
| 16.02 | 40.67 | 1.8977 | 1.3052 | 26.8406476748653 |
| 548.23 | 521.01 | 0.9942 | 1.16 | 178.903619329027 |
| 4.79 | 727.74 | 0.0523 | 2.8567 | 17.3808040274405 |
| 8.7 | 24950.27 | 0.13 | 26.1316 | 27.3596819321835 |
| 678.17 | 552.51 | 0.369 | 0.3909 | 81.0973618704188 |
| 39.93 | 44.79 | 0.0088 | 0.0079 | 56.7727884661327 |
| 32.55 | 142.33 | 0.2689 | 0.5395 | 37.8653064757609 |
| 10.96 | 5266.86 | 0.3845 | 13.6622 | 18.5093516053457 |
| 42.38 | 50.41 | 0.4296 | 0.2032 | 31.7393149214543 |
| 5.34 | 51.46 | 0.0666 | 0.7061 | 13.3172607487842 |
| 23.29 | 339.46 | 1.5186 | 3.5276 | 37.1216878447972 |
| 9.04 | 12.1 | 0.0037 | 0.0069 | 24.8729107083459 |
| 47.14 | 66.29 | 0.2647 | 0.269 | 43.7064025692658 |
| 17.94 | 6294.58 | 0.1836 | 6.2149 | 32.3793446584418 |
| 17.29 | 14.41 | 0.0059 | 0.0072 | 31.6626326861216 |
| 19.16 | 12.53 | 0.0179 | 0.0102 | 24.9543045059329 |
| 24.02 | 87.42 | 0.0793 | 0.1097 | 21.7529558099119 |
| 10.89 | 518.29 | 0.3672 | 3.4473 | 15.7483704077821 |
| 5.91 | 35.49 | 0.0043 | 0.0891 | 20.9216242244549 |
| 1437.3 | 5744.3 | 1.1903 | 1.5342 | 129.520534128152 |
| 16.07 | 30.89 | 0.1341 | 0.5892 | 16.2095593605783 |
| 4.3 | 7.36 | 0.0325 | 0.1159 | 14.1906639073741 |
| 14.38 | 10.24 | 0.1253 | 0.2286 | 20.9792856708997 |
| 34.57 | 20.31 | 3.6634 | 2.6604 | 20.4552830676368 |
| 15.54 | 10.16 | 0.5058 | 0.7833 | 19.9165680690871 |
| 28.51 | 3523.7 | 0.0039 | 0.244 | 33.8216898309303 |
| 3.36 | 3.49 | 0.1553 | 0.2156 | 12.6870832271861 |
| 22.66 | 35.47 | 3.3391 | 2.3523 | 43.8671132037225 |
| 32.86 | 36.03 | 1.007 | 1.0153 | 42.3626526400323 |
| 8.94 | 32.85 | 0.8045 | 1.6127 | 14.4606956396263 |
| 150.82 | 1200.14 | 1.3606 | 1.9972 | 61.6643818511378 |
| 85.38 | 179.47 | 0.1901 | 0.2962 | 119.087341399158 |
| 177.8 | 7166.9 | 0.4142 | 3.7847 | 80.2958492550376 |
| 9.4 | 72.53 | 0.01 | 0.0518 | 36.7230100628799 |
| 34.22 | 7851.25 | 0.1394 | 4.3146 | 61.1568482162493 |
| 236.73 | 174.01 | 0.0055 | 0.0052 | 29.8435750223415 |

| | | | | |
|---|---|---|---|---|
| 29.91 | 6762.06 | 0.0114 | 0.9909 | 33.1815406658833 |
| 9.86 | 2141.16 | 0.0942 | 7.0656 | 44.3097291741724 |
| 7.8 | 13921.86 | 0.0816 | 12.4753 | 16.2982298256813 |
| 48.9 | 122.44 | 0.0317 | 0.1055 | 33.5364241693682 |
| 28 | 120.93 | 0.0661 | 0.2905 | 29.1722907483505 |
| 27.41 | 4078.77 | 0.2283 | 5.4334 | 42.7394391824007 |
| 5.7 | 12.76 | 0.4355 | 0.6234 | 13.7807280421236 |
| 19.5 | 16.11 | 0.0084 | 0.005 | 31.5683595850688 |
| 20.45 | 42 | 0.2237 | 0.2519 | 92.1434451160565 |

| MHCflurry_BAWT | MHCflurry_Pmut | MHCflurry_PWT | mixMHCpred_Smut | mixMHCpred_SWT |
|---|---|---|---|---|
| 55.7643867214409 | 0.12225 | 0.119 | 0.459492 | 0.434462 |
| 20.2970858867476 | 0.041125 | 0.0995 | 0.503552 | 0.525125 |
| 12.8930029193755 | 0.012125 | 0.021625 | 0.484827 | 0.379703 |
| 20.9508694973094 | 0.043625 | 0.1075 | 0.467204 | 0.475231 |
| 17.864391909566 | 0.043625 | 0.0755 | 0.268879 | 0.331325 |
| 289.203004874046 | 0.317125 | 1.061 | 0.146396 | 0.017172 |
| 34.8456481081839 | 0.053 | 0.09325 | 0.355603 | 0.295237 |
| 1556.32042580725 | 0.20825 | 2.07525 | 0.277563 | -0.186112 |
| 64.8888311981951 | 0.14275 | 0.17625 | 0.466438 | 0.423216 |
| 28.2741746984835 | 0.009625 | 0.004125 | 0.750597 | 0.76968 |
| 5139.15717749382 | 3.456375 | 3.9385 | 0.013513 | -0.112982 |
| 13.6208291808157 | 0.02825 | 0.0295 | 0.54346 | 0.594395 |
| 14.6704415550145 | 0.033125 | 0.03975 | 0.236657 | 0.185851 |
| 10.9506902620108 | 0.002875 | 0.00575 | 0.579498 | 0.451986 |
| 21.6392095067484 | 0.03675 | 0.115 | 0.211839 | 0.226468 |
| 116.673652518787 | 0.14325 | 0.395125 | 0.480404 | 0.318127 |
| 51.5709499767421 | 0.1645 | 0.365375 | -0.002622 | -0.139183 |
| 214.503898827464 | 0.843624999999999 | 0.923499999999999 | -0.092958 | -0.159418 |
| 228.4040612218 | 0.0695 | 0.952124999999999 | 0.207173 | -0.324011 |
| 15468.5245179948 | 0.168875 | 8.213375 | 0.278502 | -0.27399 |
| 85.6274033457948 | 0.286125 | 0.30625 | 0.123865 | 0.108155 |
| 56.1847844565313 | 0.016375 | 0.013 | 0.668858 | 0.650254 |
| 102.105440281088 | 0.264125 | 0.617625 | 0.151624 | 0.129235 |
| 6543.79829476495 | 0.081 | 4.093 | 0.11633 | -0.345337 |
| 26.8887565252625 | 0.210875 | 0.1645 | 0.155392 | 0.174605 |
| 39.0004667425853 | 0.027125 | 0.274375 | 0.42159 | 0.235796 |
| 248.701009704952 | 0.2575 | 0.991624999999999 | 0.121092 | 0.008734 |
| 23.7272516972457 | 0.005875 | 0.003875 | 0.587022 | 0.570937 |
| 50.9653137435918 | 0.309625 | 0.358625 | 0.038149 | 0.008831 |
| 5905.0391493356 | 0.042875 | 2.8935 | 0.191324 | -0.276443 |
| 32.6127942561405 | 0.038125 | 0.042875 | 0.516126 | 0.523281 |
| 21.9995772676232 | 0.150625 | 0.1195 | 0.59886 | 0.633968 |
| 34.4147188897565 | 0.116875 | 0.231875 | 0.496712 | 0.417114 |
| 146.928287076206 | 0.0505 | 0.757499999999999 | 0.066163 | -0.284714 |
| 36.1301900274018 | 0.0015 | 0.106625 | 0.693252 | 0.522204 |
| 143.595526309384 | 0.706499999999999 | 0.747749999999999 | 0.119356 | 0.140929 |
| 21.6008733477579 | 0.055875 | 0.115 | 0.127589 | 0.144495 |
| 17.7602790318868 | 0.035375 | 0.073625 | 0.50577 | 0.401005 |
| 18.2761664221088 | 0.110375 | 0.079125 | 0.259484 | 0.2735 |
| 25.0854976062881 | 0.10225 | 0.150625 | -0.483549 | -0.406748 |
| 17.4579043973462 | 0.097 | 0.07125 | 0.165021 | 0.244619 |
| 204.217371881419 | 0.010375 | 0.428125 | 0.806259 | 0.450631 |
| 13.363362279342 | 0.019375 | 0.027125 | 0.096373 | 0.068867 |
| 79.6703197808179 | 0.3145 | 0.516749999999999 | 0.01414 | -0.0802 |
| 39.4936127809678 | 0.29925 | 0.278125 | 0.303375 | 0.333659 |
| 21.2445648529769 | 0.03825 | 0.11325 | 0.352182 | 0.360103 |
| 109.068691484341 | 0.424875 | 0.641875 | -0.177138 | -0.155564 |
| 128.17630195003 | 0.168 | 0.182125 | 0.501002 | 0.445194 |
| 5725.22519952387 | 0.516749999999999 | 3.775 | 0.208084 | -0.255591 |
| 49.3211092956147 | 0.02425 | 0.097625 | 0.437252 | 0.284034 |
| 18052.4209112818 | 0.170875 | 6.603625 | 0.293954 | -0.026215 |
| 29.1502045693032 | 0.001 | 0.0005 | 0.603259 | 0.589903 |

| | | | | |
|---|---|---|---|---|
| 105.767442081471 | 0.00975 | 0.361 | 0.540642 | 0.140434 |
| 3974.00423115778 | 0.1045 | 4.27825 | 0.465079 | -0.125658 |
| 7722.34544612093 | 0.055875 | 4.5225 | 0.536668 | -0.015824 |
| 39.4225752855077 | 0.03825 | 0.0635 | 0.352772 | 0.21748 |
| 60.312116982767 | 0.185875 | 0.417 | 0.380484 | 0.180361 |
| 5184.0003012715 | 0.303375 | 3.575875 | 0.277338 | -0.353067 |
| 18.398109932266 | 0.03125 | 0.079125 | -0.144871 | -0.163358 |
| 28.426176076095 | 0.00425 | 0.001625 | 0.805123 | 0.84195 |
| 114.818167263705 | 0.124375 | 0.160125 | 0.546258 | 0.404212 |

| mixMHCpred_Rmut | mixMHCpred_RWT | deepimmunemut | deepimmuneWT | CIImmmut | CIImmWT |
|---|---|---|---|---|---|
| 0.04 | 0.05 | 0.9888036 | 0.9885606 | 0.17422 | 0.20212 |
| 0.02 | 0.02 | 0.7304131 | 0.81391066 | 0.13176 | 0.15282 |
| 0.03 | 0.07 | 0.8593644 | 0.86599874 | 0.11016 | 0.08478 |
| 0.03 | 0.03 | 0.9262525 | 0.9153544 | 0.20052 | -0.07458 |
| 0.2 | 0.1 | 0.87382555 | 0.83095276 | -0.21999 | -0.18489 |
| 0.4 | 0.9 | 0.5311489 | 0.5254253 | 0.1914 | 0.16257 |
| 0.05 | 0.08 | 0.5224895 | 0.7445742 | -0.06846 | 0.23904 |
| 0.2 | 4 | 0.93755025 | 0.9295901 | 0.18454 | 0.18454 |
| 0.04 | 0.06 | 0.6093736 | 0.62421256 | -0.16194 | -0.21084 |
| 0.01 | 0.01 | 0.3877036 | 0.34272885 | -0.28164 | 0.03611 |
| 1 | 3 | 0.22146109 | 0.19294548 | -0.26673 | -0.10472 |
| 0.02 | 0.01 | 0.8222445 | 0.76088524 | 0.16095 | 0.16401 |
| 0.2 | 0.3 | 0.6299958 | 0.6770478 | -0.01226 | -0.04286 |
| 0.01 | 0.04 | 0.83397555 | 0.81340957 | 0.03405 | 0.03405 |
| 0.3 | 0.3 | 0.64573586 | 0.63626593 | 0.16314 | 0.16314 |
| 0.03 | 0.1 | 0.92794204 | 0.92212766 | -0.1513 | -0.13332 |
| 2 | 3 | 0.84704894 | 0.7750858 | -0.15568 | -0.05621 |
| 2 | 4 | 0.82352835 | 0.8992623 | 0.27171 | 0.01119 |
| 0.3 | 10 | 0.7739066 | 0.7251859 | 0.20938 | 0.20938 |
| 0.2 | 7 | 0.82194304 | 0.61427975 | 0.23374 | 0.23374 |
| 0.4 | 0.5 | 0.79586554 | 0.8945234 | -0.04792 | -0.04762 |
| 0.01 | 0.01 | 0.8794873 | 0.85953116 | 0.17141 | -0.09452 |
| 0.4 | 0.5 | 0.7574198 | 0.725669 | 0.12444 | 0.03604 |
| 0.5 | 11 | 0.55906916 | 0.6176267 | 0.16191 | 0.16191 |
| 0.4 | 0.4 | 0.6210585 | 0.76889044 | -0.10143 | -0.10169 |
| 0.05 | 0.2 | 0.6809099 | 0.7255723 | 0.08519 | 0.08519 |
| 0.5 | 1 | 0.56727153 | 0.5178597 | 0.12163 | 0.12163 |
| 0.01 | 0.01 | 0.8452617 | 0.83429503 | 0.03417 | 0.03417 |
| 0.8 | 1 | 0.8896481 | 0.86354554 | -0.02657 | -0.23706 |
| 0.3 | 9 | 0.94178665 | 0.8625609 | 0.31858 | 0.31858 |
| 0.01 | 0.01 | 0.9700553 | 0.954083 | 0.04304 | -0.24754 |
| 0.01 | 0.01 | 0.68797344 | 0.7674557 | -0.26671 | -0.20839 |
| 0.03 | 0.05 | 0.9119426 | 0.91801107 | -0.11603 | -0.11603 |
| 0.7 | 8 | 0.86094743 | 0.8294928 | 0.22151 | 0.22151 |
| 0.01 | 0.01 | 0.7872757 | 0.73260206 | -0.05566 | -0.05566 |
| 0.5 | 0.4 | 0.5414773 | 0.6485951 | 0.27298 | 0.29404 |
| 0.5 | 0.4 | 0.77423406 | 0.64705884 | 0.29214 | 0.12564 |
| 0.02 | 0.06 | 0.850978 | 0.8112042 | 0.20056 | 0.21854 |
| 0.2 | 0.2 | 0.87241477 | 0.8432344 | -0.1502 | -0.10322 |
| 25 | 16 | 0.5365665 | 0.67431724 | 0.41234 | 0.14641 |
| 0.4 | 0.2 | 0.5247706 | 0.71047586 | 0.08716 | 0.08716 |
| 0.01 | 0.05 | 0.77572906 | 0.75044507 | 0.03205 | 0.03205 |
| 0.6 | 0.7 | 0.8619778 | 0.7722631 | 0.08083 | 0.24403 |
| 1 | 2 | 0.8573376 | 0.68062 | -0.23622 | -0.12989 |
| 0.2 | 0.1 | 0.7057327 | 0.5843926 | -0.31661 | -0.21617 |
| 0.08 | 0.08 | 0.7172195 | 0.69647515 | -0.10348 | -0.19528 |
| 4 | 3 | 0.300511 | 0.47679156 | 0.00769 | 0.02875 |
| 0.02 | 0.02 | 0.31533828 | 0.47016972 | 0.09841 | 0.09831 |
| 0.3 | 6 | 0.60148644 | 0.5589822 | 0.28774 | 0.28774 |
| 0.05 | 0.2 | 0.64054286 | 0.33626765 | 0.01873 | 0.01873 |
| 0.2 | 2 | 0.8002486 | 0.80364954 | -0.03854 | -0.03854 |
| 0.02 | 0.02 | 0.73621327 | 0.710712 | 0.40786 | 0.31912 |

| | | | | | |
|---|---|---|---|---|---|
| 0.02 | 0.4 | 0.57582885 | 0.52368987 | -0.11382 | -0.11382 |
| 0.04 | 3 | 0.7339178 | 0.7806962 | 0.17099 | 0.17099 |
| 0.02 | 2 | 0.9579121 | 0.9272148 | -0.16757 | -0.16757 |
| 0.05 | 0.2 | 0.7002374 | 0.7328023 | -0.34867 | -0.18045 |
| 0.07 | 0.3 | 0.7836723 | 0.60934466 | 0.09312 | 0.09942 |
| 0.2 | 12 | 0.67874706 | 0.65457356 | 0.0909 | 0.0909 |
| 3 | 4 | 0.8723961 | 0.84512603 | 0.01322 | 0.06662 |
| 0.01 | 0.01 | 0.41928127 | 0.34251702 | 0.38329 | 0.22009 |
| 0.01 | 0.03 | 0.41808566 | 0.32515496 | 0.00583 | -0.03647 |

| deepitopemut | deepitopeWT | PRIMEmut | PRIMEwt | netMHC_I.DAIBA | netMHC_I.DAIR |
|---|---|---|---|---|---|
| 1 | 1 | 0.199138 | 0.194789 | 23.79 | -0.001 |
| 1 | 1 | 0.208258 | 0.199386 | 77.44 | 0.0656 |
| 1 | 1 | 0.198485 | 0.199187 | -0.4 | 0.0081 |
| 1 | 1 | 0.210145 | 0.196929 | 7.66 | 0.0041 |
| 1 | 1 | 0.193517 | 0.190149 | 24.48 | -0.0391 |
| 1 | 1 | 0.176441 | 0.170908 | 649.05 | 1.44 |
| 0 | 0 | 0.196345 | 0.19089 | 6.95 | 0.1081 |
| 0 | 0 | 0.197441 | 0.175838 | 2003.17 | 4.5067 |
| 0 | 0 | 0.194652 | 0.187535 | 65.42 | 0.186 |
| 1 | 1 | 0.187992 | 0.185601 | -237.22 | -0.0044 |
| 1 | 1 | 0.154484 | 0.146376 | 3197.23 | 2.5683 |
| 1 | 1 | 0.195913 | 0.203921 | -0.21 | -0.0105 |
| 1 | 1 | 0.184972 | 0.182083 | 0.41 | 0.0269 |
| 1 | 1 | 0.210814 | 0.19211 | 0.77 | -0.006 |
| 1 | 1 | 0.187346 | 0.187346 | 29.97 | 0.0725 |
| 0 | 0 | 0.196235 | 0.185651 | 11.48 | 0.0516 |
| 1 | 1 | 0.180155 | 0.171087 | 24.65 | -0.5925 |
| 0 | 0 | 0.173592 | 0.167215 | -27.22 | 0.1658 |
| 1 | 1 | 0.191886 | 0 | 722.95 | 2.8044 |
| 1 | 0 | 0.185455 | 0 | 24941.57 | 26.0016 |
| 0 | 0 | 0.178478 | 0.173429 | -125.66 | 0.0219 |
| 1 | 1 | 0.214718 | 0.203923 | 4.86 | -0.0009 |
| 1 | 1 | 0.174791 | 0.170139 | 109.78 | 0.2706 |
| 1 | 1 | 0.187861 | 0 | 5255.9 | 13.2777 |
| 1 | 1 | 0.175556 | 0.17998 | 8.02999999999999 | -0.2264 |
| 1 | 1 | 0.195384 | 0.18523 | 46.12 | 0.6395 |
| 0 | 0 | 0.18309 | 0.178212 | 316.17 | 2.009 |
| 1 | 1 | 0.203124 | 0.196673 | 3.06 | 0.0032 |
| 0 | 0 | 0.175516 | 0.173986 | 19.15 | 0.0043 |
| 1 | 1 | 0.190932 | 0 | 6276.64 | 6.0313 |
| 1 | 1 | 0.213499 | 0.211871 | -2.88 | 0.0013 |
| 1 | 1 | 0.187575 | 0.187314 | -6.63 | -0.0077 |
| 1 | 1 | 0.202531 | 0.190733 | 63.4 | 0.0304 |
| 1 | 1 | 0.192697 | 0 | 507.4 | 3.0801 |
| 1 | 1 | 0.207908 | 0.207908 | 29.58 | 0.0848 |
| 1 | 1 | 0.175957 | 0.169658 | 4307 | 0.3439 |
| 1 | 0 | 0.188704 | 0.183335 | 14.82 | 0.4551 |
| 1 | 1 | 0.214304 | 0.20564 | 3.06 | 0.0834 |
| 1 | 1 | 0.178102 | 0.184794 | -4.14 | 0.1033 |
| 1 | 0 | 0 | 0 | -14.26 | -1.003 |
| 0 | 1 | 0.182342 | 0.195104 | -5.38 | 0.2775 |
| 1 | 1 | 0.196913 | 0.185092 | 3495.19 | 0.2401 |
| 1 | 1 | 0.17614 | 0.175885 | 0.13 | 0.0603 |
| 0 | 0 | 0.168302 | 0.163739 | 12.81 | -0.9868 |
| 0 | 0 | 0.184566 | 0.189578 | 3.17 | 0.0083 |
| 0 | 0 | 0.197253 | 0.196805 | 23.91 | 0.8082 |
| 1 | 1 | 0.16349 | 0.157959 | 1049.32 | 0.6366 |
| 0 | 0 | 0.198493 | 0.198493 | 94.09 | 0.1061 |
| 1 | 1 | 0.197472 | 0 | 6989.1 | 3.3705 |
| 1 | 1 | 0.181242 | 0.171665 | 63.13 | 0.0418 |
| 1 | 0 | 0.183374 | 0.167544 | 7817.03 | 4.1752 |
| 1 | 1 | 0.204772 | 0.204404 | -62.72 | -0.0003 |

| | | | | | |
|---|---|---|---|---|---|
| 1 | 0 | 0.19073 | 0.169706 | 6732.15 | 0.9795 |
| 1 | 1 | 0.187275 | 0.158011 | 2131.3 | 6.9714 |
| 0 | 0 | 0.187953 | 0.156788 | 13914.06 | 12.3937 |
| 1 | 1 | 0.181736 | 0.172847 | 73.54 | 0.0738 |
| 1 | 1 | 0.188087 | 0.177747 | 92.93 | 0.2244 |
| 1 | 0 | 0.186275 | 0 | 4051.36 | 5.2051 |
| 1 | 1 | 0.170945 | 0.169015 | 7.06 | 0.1879 |
| 1 | 1 | 0.18619 | 0.185516 | -3.39 | -0.0034 |
| 1 | 1 | 0.208465 | 0.199245 | 21.55 | 0.0282 |

| netMHC_II.DAIBA | netMHC_II.DAIR | flurry_I.DAIBA | flurry_I.DAIP |
|---|---|---|---|
| 0.77840909 | 1.05617978 | -0.7716245 | -0.00325 |
| 0.187152303978167 | 0.468825910931174 | 5.47249171240341 | 0.058375 |
| 1.03613369467028 | 0.487341772151899 | 1.21332825862091 | 0.0095 |
| 0.512722646310433 | 0.840466926070039 | 5.94375971573642 | 0.063875 |
| 0.364980544747082 | 1.40644490644491 | 2.71184118519613 | 0.031875 |
| 0.294526205952045 | 0.263050153531218 | 244.742390916134 | 0.743875 |
| 0.756140350877193 | 0.720455133178174 | 5.27433594585155 | 0.04025 |
| 0.008454399208019 | 0.094640202499096 | 1524.98132598497 | 1.867 |
| 0.72687040748163 | 0.470236399886072 | 6.9306934101107 | 0.0335 |
| 1.84398904187569 | 1.88 | -6.62228202342268 | -0.0055 |
| 0.641242779366155 | 0.692625305184547 | 1314.76847417997 | 0.482124999999999 |
| 1.07664233576642 | 2 | 0.149945757373317 | 0.00125 |
| 0.895939086294416 | 0.820666666666667 | 0.684479874624971 | 0.006625 |
| 0.798429319371728 | 2.2 | 0.391167606313889 | 0.002875 |
| 0.188903924221922 | 0.439288476411446 | 7.3273033652573 | 0.07825 |
| 0.801966534414352 | 0.659405940594059 | 48.6105432804363 | 0.251875 |
| 0.393902139168921 | 1.45395341710083 | 24.7303023018767 | 0.200875 |
| 1.05224467860502 | 0.857068965517241 | 35.6002794984372 | 0.079875 |
| 0.006582021051474 | 0.018307837714846 | 211.023257194359 | 0.882624999999999 |
| 0.000348693621352 | 0.004974819758453 | 15441.1648360626 | 8.0445 |
| 1.22743479756023 | 0.943975441289332 | 4.53004147537602 | 0.020125 |
| 0.891493636972538 | 1.11392405063291 | -0.58800400960142 | -0.003375 |
| 0.228693880418745 | 0.498424467099166 | 64.2401338053271 | 0.3535 |
| 0.002080936269428 | 0.028143344410124 | 6525.2889431596 | 4.012 |
| 0.840706209085499 | 2.11417322834646 | -4.85055839619181 | -0.046375 |
| 0.10376991838321 | 0.094320917717037 | 25.6832059938011 | 0.24725 |
| 0.068608967183173 | 0.430490985372491 | 211.579321860155 | 0.734124999999999 |
| 0.747107438016529 | 0.536231884057971 | -1.14565901110015 | -0.002 |
| 0.71111781565847 | 0.984014869888476 | 7.25891117432598 | 0.049 |
| 0.002850071013475 | 0.029541907351687 | 5872.65980467716 | 2.850625 |
| 1.1998612074948 | 0.819444444444444 | 0.950161570018921 | 0.00475 |
| 1.52913008778931 | 1.75490196078431 | -2.95472723830971 | -0.031125 |
| 0.27476549988561 | 0.722880583409298 | 12.6617630798446 | 0.115 |
| 0.021011402882556 | 0.106518144634932 | 131.179916668424 | 0.706999999999999 |
| 0.166525781910397 | 0.048260381593715 | 15.208565802947 | 0.105125 |
| 0.250213254878749 | 0.775844088124104 | 14.074992181232 | 0.04125 |
| 0.520233085140822 | 0.227596741344196 | 5.39131398717952 | 0.059125 |
| 0.584239130434783 | 0.280414150129422 | 3.56961512451276 | 0.03825 |
| 1.404296875 | 0.548118985126859 | -2.70311924879092 | -0.03125 |
| 1.70211718365337 | 1.37701097579311 | 4.63021453865136 | 0.048375 |
| 1.52952755905512 | 0.645729605515128 | -2.45866367174094 | -0.02575 |
| 0.008090927150438 | 0.015983606557377 | 170.395682050488 | 0.41775 |
| 0.962750716332378 | 0.720315398886827 | 0.676279052155921 | 0.00775 |
| 0.638849732168029 | 1.41950431492582 | 35.8032065770953 | 0.20225 |
| 0.912017762975298 | 0.991825076332118 | -2.8690398590645 | -0.021125 |
| 0.272146118721461 | 0.498852855459788 | 6.78386921335051 | 0.075 |
| 0.125668671988268 | 0.68125375525736 | 47.4043096332032 | 0.217 |
| 0.475734105978715 | 0.641796083727211 | 9.08896055087199 | 0.014125 |
| 0.024808494607152 | 0.109440642587259 | 5644.92935026884 | 3.25825 |
| 0.129601544188612 | 0.193050193050193 | 12.5980992327348 | 0.073375 |
| 0.004358541633498 | 0.03230890464933 | 17991.2640630655 | 6.43275 |
| 1.36043905522671 | 1.05769230769231 | -0.693370453038295 | -0.0005 |

| | | | |
|---|---|---|---|
| 0.00442320831226 | 0.011504692703603 | 72.5859014155876 | 0.35125 |
| 0.004604980477872 | 0.013332201086957 | 3929.6945019836 | 4.17375 |
| 0.000560269963927 | 0.006540924867538 | 7706.04721629525 | 4.466625 |
| 0.39937928781444 | 0.300473933649289 | 5.88615111613951 | 0.02525 |
| 0.23153890680559 | 0.227538726333907 | 31.1398262344165 | 0.231125 |
| 0.006720163186451 | 0.042017889351051 | 5141.2608620891 | 3.2725 |
| 0.446708463949843 | 0.698588386268848 | 4.61738189014241 | 0.047875 |
| 1.21042830540037 | 1.68 | -3.14218350897379 | -0.002625 |
| 0.486904761904762 | 0.888050813815006 | 22.6747221476488 | 0.03575 |

| flurry_II.DAIBA | flurry_II.DAIP |
|---|---|
| 1.01383722 | 1.02731092 |
| 0.730380422936648 | 0.413316582914573 |
| 0.905892501055938 | 0.560693641618497 |
| 0.716300093583241 | 0.405813953488372 |
| 0.848198516975884 | 0.577814569536424 |
| 0.153734965434664 | 0.298892554194156 |
| 0.848637168995206 | 0.568364611260054 |
| 0.020136662927892 | 0.100349355499337 |
| 0.893191273719483 | 0.809929078014184 |
| 1.23421663387324 | 2.33333333333333 |
| 0.744166518210846 | 0.877586644661673 |
| 0.988991436910133 | 0.957627118644068 |
| 0.953342926178591 | 0.833333333333333 |
| 0.964279182685781 | 0.5 |
| 0.661387660072693 | 0.319565217391304 |
| 0.583363148139987 | 0.362543498892756 |
| 0.520460602082571 | 0.450222374273008 |
| 0.834034347659703 | 0.913508391987006 |
| 0.07609673809855 | 0.0729946173034 |
| 0.001768732492899 | 0.020560975238559 |
| 0.947095891053918 | 0.934285714285714 |
| 1.01046553822871 | 1.25961538461538 |
| 0.370845141762484 | 0.427646225460433 |
| 0.002828533333638 | 0.019789885169802 |
| 1.18039355563485 | 1.28191489361702 |
| 0.341464137767429 | 0.098861047835991 |
| 0.149262312560921 | 0.259674776251103 |
| 1.04828452219071 | 1.51612903225806 |
| 0.857571539521059 | 0.86336702683862 |
| 0.005483341234424 | 0.014817694833247 |
| 0.970865373799118 | 0.889212827988338 |
| 1.13430836430926 | 1.26046025104603 |
| 0.63208291427848 | 0.504043126684637 |
| 0.107184060477163 | 0.066666666666667 |
| 0.579062114220476 | 0.014067995310668 |
| 0.901981680467491 | 0.944834503510532 |
| 0.750412221747547 | 0.485869565217391 |
| 0.799011315188018 | 0.480475382003396 |
| 1.14790406184532 | 1.39494470774092 |
| 0.815422655299821 | 0.678838174273859 |
| 1.14083383754322 | 1.36140350877193 |
| 0.165616125206867 | 0.024233576642336 |
| 0.949393046598657 | 0.714285714285714 |
| 0.550607972007718 | 0.608611514271892 |
| 1.07264566741403 | 1.07595505617978 |
| 0.680677422187824 | 0.337748344370861 |
| 0.565371978080355 | 0.661927945472249 |
| 0.92909016399603 | 0.922443376801647 |
| 0.014024924165728 | 0.136887417218543 |
| 0.744569832011971 | 0.248399487836108 |
| 0.003387736665171 | 0.025875939351493 |
| 1.02378612648806 | 2 |

| | |
|---|---|
| 0.313721689897011 | 0.027008310249307 |
| 0.011149894815604 | 0.02442587506574 |
| 0.002110528457888 | 0.012354892205639 |
| 0.850690852296924 | 0.602362204724409 |
| 0.483688721400476 | 0.44574340527578 |
| 0.008244490103891 | 0.084839374978152 |
| 0.749029552103905 | 0.394944707740916 |
| 1.11053838196746 | 2.61538461538462 |
| 0.802516250798785 | 0.776736924277909 |
