## Supplementary Table 2 for "MHC-I binding affinity derived metrics fail to predict tumor specific neoantigen immunogenicity"

**Validation peptides – negative set**

| Peptide | HLA | netMHCp | netMH( | mixMHCpre( | mixMHCpre | CIImm | deepitope |
|---|---|---|---|---|---|---|---|
| FLLGKTSSV | HLA-A02:01 | 3.51 | 0.006 | 0.549536 | 0.02 | -0.37924 | 1 |
| SLLSLLYAL | HLA-A02:01 | 4.25 | 0.079 | 0.599057 | 0.01 | -0.17157 | 1 |
| FQLNQSFEI | HLA-A02:01 | 7.28 | 0.21 | 0.157263 | 0.4 | -0.12134 | 1 |
| FILDAVQRV | HLA-A02:01 | 6.42 | 0.01 | 0.547807 | 0.02 | 0.02816 | 1 |
| HLFCVFYGL | HLA-A02:01 | 11.74 | 1.136 | 0.264974 | 0.2 | 0.15083 | 1 |
| SLAPLSPRV | HLA-A02:01 | 8.09 | 0.006 | 0.585183 | 0.01 | -0.14411 | 1 |
| RMLDKNPEV | HLA-A02:01 | 6.29 | 0.011 | 0.735191 | 0.01 | -0.14823 | 1 |
| HLMDGDLGL | HLA-A02:01 | 8.68 | 0.035 | 0.378374 | 0.07 | 0.02964 | 1 |
| LLGDPGWRV | HLA-A02:01 | 20.57 | 0.07 | 0.077854 | 0.6 | 0.2716 | 1 |
| FSFSLDFLV | HLA-A02:01 | 12.37 | 1.132 | -0.268254 | 7 | -0.02607 | 1 |
| MLFLRFCYI | HLA-A02:01 | 11.58 | 1.557 | -0.168278 | 4 | 0.13978 | 1 |
| FLQKYTVKL | HLA-A02:01 | 8.3 | 0.018 | 0.52844 | 0.02 | -0.31282 | 1 |
| CLFLEIYTV | HLA-A02:01 | 11.08 | 0.435 | 0.340121 | 0.09 | 0.26918 | 0 |
| FLEIYTVTV | HLA-A02:01 | 8.12 | 0.108 | 0.166985 | 0.4 | 0.25688 | 1 |
| SLSTSLSSV | HLA-A02:01 | 16.4 | 0.087 | 0.452798 | 0.04 | -0.42246 | 1 |
| SVVDVFFQL | HLA-A02:01 | 30.06 | 0.103 | 0.23023 | 0.3 | 0.21724 | 1 |
| GLNETIAKL | HLA-A02:01 | 35.16 | 0.026 | 0.600438 | 0.01 | 0.16875 | 1 |
| YTAPYPHPA | HLA-A02:01 | 60.83 | 0.46 | -0.137951 | 3 | 0.00832 | 1 |
| ALVGAISSI | HLA-A02:01 | 38.61 | 0.229 | 0.240371 | 0.2 | -0.0254 | 1 |
| SLSSVTLLL | HLA-A02:01 | 18.93 | 0.107 | 0.364191 | 0.08 | -0.15927 | 1 |
| YTSEHAASV | HLA-A02:01 | 37.04 | 0.246 | 0.300043 | 0.2 | 0.05174 | 1 |
| AICKPLHYV | HLA-A02:01 | 138.05 | 0.361 | 0.205782 | 0.3 | -0.2306 | 0 |
| ALLMCLLPL | HLA-A02:01 | 25.45 | 1.964 | -0.007715 | 2 | -0.25908 | 1 |
| YLDELIRNT | HLA-A02:01 | 71.23 | 0.155 | 0.199927 | 0.3 | 0.26233 | 1 |
| WMSDSGTRL | HLA-A02:01 | 176.33 | 0.742 | 0.149528 | 0.4 | -0.09758 | 1 |
| KLLPTNTNI | HLA-A02:01 | 35.58 | 0.061 | 0.431214 | 0.04 | 0.04593 | 1 |
| SLDLPLNPL | HLA-A02:01 | 154.98 | 0.479 | -0.133836 | 3 | -0.03714 | 1 |
| CMQANSHYA | HLA-A02:01 | 203.32 | 2.984 | 0.154171 | 0.4 | -0.13512 | 0 |
| FSDYYDLSY | HLA-A01:01 | 2.35 | 0.002 | 0.815317 | 0.01 | -0.08526 | 1 |
| YSSALDLCY | HLA-A01:01 | 7.13 | 0.04 | 0.487766 | 0.02 | -0.04511 | 1 |
| FSDKKVGTY | HLA-A01:01 | 11.72 | 0.004 | 0.72709 | 0.01 | -0.32966 | 1 |
| HSEYSSFFY | HLA-A01:01 | 6.52 | 0.01 | 0.632535 | 0.01 | -0.12085 | 1 |
| CSNFLLLAY | HLA-A01:01 | 74.55 | 0.465 | 0.423455 | 0.03 | 0.10796 | 0 |
| FTGTISVMY | HLA-A01:01 | 34.81 | 0.065 | 0.323939 | 0.07 | -0.04383 | 1 |
| FTGTISVMY | HLA-A26:01 | 804.35 | 0.195 | -0.105252 | 3 | -0.04383 | 1 |
| QTQSVVFLY | HLA-A01:01 | 68.75 | 0.064 | 0.456973 | 0.02 | -0.03269 | 1 |
| CTDTYMLEL | HLA-A01:01 | 95.53 | 0.517 | 0.423397 | 0.03 | -0.0735 | 0 |
| SSDSQEENY | HLA-A01:01 | 34.47 | 0.007 | 0.702937 | 0.01 | -0.0971 | 1 |
| LTSMAYDCY | HLA-A01:01 | 21.38 | 0.153 | 0.323685 | 0.07 | -0.20856 | 1 |
| WADWGHRTY | HLA-A01:01 | 111.63 | 0.097 | 0.376263 | 0.04 | 0.3599 | 1 |
| VSDGFTAVM | HLA-A01:01 | 296.83 | 0.403 | 0.157093 | 0.3 | 0.24898 | 1 |
| WSCLGHLGY | HLA-A01:01 | 113.34 | 0.418 | 0.472118 | 0.02 | 0.04523 | 0 |
| YTFLIFSDY | HLA-A01:01 | 255.93 | 0.394 | 0.317134 | 0.07 | 0.13998 | 1 |
| YTFLIFSDY | HLA-A26:01 | 130.48 | 0.133 | 0.211032 | 0.2 | 0.13998 | 1 |
| LSFFTVMAY | HLA-A01:01 | 240.19 | 0.487 | 0.194275 | 0.2 | 0.10712 | 1 |
| GTFFVLAYY | HLA-A01:01 | 952.21 | 0.673 | 0.194668 | 0.2 | 0.21642 | 1 |
| GTFFVLAYY | HLA-A26:01 | 599.2 | 0.252 | 0.102279 | 0.5 | 0.21642 | 1 |
| LVVLAPLLY | HLA-A01:01 | 1032.09 | 0.537 | 0.132816 | 0.4 | 0.01406 | 1 |
| YSSEENLIF | HLA-A01:01 | 931.29 | 0.551 | 0.18332 | 0.3 | 0.20686 | 1 |
| SSYIASFTY | HLA-A01:01 | 427.23 | 0.248 | 0.179297 | 0.3 | 0.13657 | 1 |
| SISSEQTFY | HLA-A01:01 | 661.79 | 0.173 | 0.296472 | 0.08 | -0.13055 | 1 |
| LTDTSLTAL | HLA-A01:01 | 889.61 | 0.594 | 0.196413 | 0.2 | -0.06966 | 1 |

| | | | | | | | |
|---|---|---|---|---|---|---|---|
| YVDVTYNFI | HLA-A01:01 | 658.06 | 1.285 | -0.052827 | 2 | 0.146 | 1 |
| HLSHQAHRY | HLA-A01:01 | 930.79 | 0.205 | 0.369816 | 0.05 | -0.03958 | 1 |
| FSDYERAEW | HLA-A01:01 | 1408.88 | 0.476 | 0.253888 | 0.2 | 0.24122 | 1 |
| VVNPIIYFY | HLA-A01:01 | 587.86 | 0.17 | 0.140648 | 0.4 | 0.3069 | 1 |
| FGYEHWALY | HLA-A01:01 | 1121.47 | 0.678 | 0.099071 | 0.5 | 0.3661 | 1 |
| FGYEHWALY | HLA-A26:01 | 527.58 | 0.404 | 0.043445 | 0.7 | 0.3661 | 1 |
| EVMKKLPLF | HLA-A26:01 | 23.08 | 0.01 | 0.806295 | 0.01 | -0.51028 | 1 |
| ETADTKVHF | HLA-A26:01 | 115.05 | 0.007 | 0.486147 | 0.02 | -0.07644 | 1 |
| FSKEDLAAM | HLA-A26:01 | 176.14 | 0.307 | 0.148159 | 0.4 | 0.09779 | 1 |
| YTIEEFMEL | HLA-A26:01 | 86.26 | 0.089 | 0.327079 | 0.07 | 0.26195 | 1 |
| ETLSPTPSY | HLA-A26:01 | 59.36 | 0.004 | 0.578945 | 0.01 | -0.25035 | 1 |
| EVSDGFTAV | HLA-A26:01 | 117.71 | 0.103 | 0.473899 | 0.02 | 0.16744 | 1 |
| ESFFWLEEM | HLA-A26:01 | 169.09 | 0.289 | 0.127139 | 0.4 | 0.50406 | 0 |
| HLIAIFCAY | HLA-A26:01 | 291.27 | 0.375 | 0.409191 | 0.04 | 0.29973 | 1 |
| YLIGVIGNF | HLA-A26:01 | 171.93 | 0.215 | 0.342498 | 0.06 | 0.2676 | 1 |
| MVILYVVYY | HLA-A26:01 | 327.72 | 0.13 | 0.336723 | 0.07 | 0.09998 | 1 |
| NTASSSLAF | HLA-A26:01 | 214.17 | 0.099 | 0.205722 | 0.2 | -0.4571 | 1 |
| EATNPYMVY | HLA-A26:01 | 222.27 | 0.024 | 0.192216 | 0.3 | -0.13227 | 1 |
| SVMYQGNTY | HLA-A26:01 | 203.13 | 0.058 | 0.209015 | 0.2 | -0.1244 | 1 |
| EIKKDGVGF | HLA-A26:01 | 363.14 | 0.083 | 0.345188 | 0.06 | -0.17886 | 1 |
| FAHFHSGEM | HLA-A26:01 | 1026.02 | 1.129 | -0.222072 | 7 | 0.09117 | 1 |
| FLFINTSSY | HLA-A26:01 | 454.5 | 0.191 | 0.123343 | 0.4 | -0.03412 | 1 |
| ESVINQGPM | HLA-A26:01 | 695.76 | 1.422 | -0.168098 | 4 | 0.0541 | 0 |
| ESVTSCEAY | HLA-A26:01 | 358.68 | 0.232 | 0.398045 | 0.04 | -0.05203 | 1 |
| YVKLANLSY | HLA-A26:01 | 443.18 | 0.091 | 0.223367 | 0.2 | -0.15517 | 1 |
| YVVTISSLL | HLA-A26:01 | 433.38 | 0.353 | 0.241253 | 0.2 | -0.11977 | 1 |
| TTATISGLV | HLA-A26:01 | 1248.1 | 1.636 | -0.070895 | 2 | 0.04775 | 1 |
| HMSSYIASF | HLA-A26:01 | 1185.41 | 0.522 | 0.356462 | 0.06 | -0.16213 | 1 |
| EVLVEYSFF | HLA-A26:01 | 401.03 | 0.199 | 0.385958 | 0.05 | 0.06074 | 1 |
| TTTATISGL | HLA-A26:01 | 1904.73 | 1.05 | 0.212944 | 0.2 | 0.09523 | 1 |
| YMAELFSFI | HLA-A26:01 | 2714.58 | 3.592 | -0.060555 | 2 | 0.14163 | 1 |
| HMIVISIAY | HLA-A26:01 | 3113.42 | 0.501 | 0.059452 | 0.7 | 0.19379 | 1 |
| LTERANFQY | HLA-A01:01 | 12.99 | 0.009 | 0.69455 | 0.01 | 0.14771 | 1 |
| FTNFWGLLI | HLA-A01:01 | 514.33 | 2.348 | -0.167369 | 4 | 0.34746 | 1 |
| FSARVLKSY | HLA-A01:01 | 627.76 | 0.269 | 0.345068 | 0.06 | -0.18412 | 1 |
| NTLEQTVPY | HLA-A01:01 | 533.71 | 0.245 | 0.18231 | 0.3 | 0.04925 | 1 |
| LVEQALRFY | HLA-A01:01 | 825.06 | 0.214 | 0.366554 | 0.05 | 0.05568 | 1 |
| SCDSLCGGY | HLA-A01:01 | 612.78 | 0.398 | 0.445001 | 0.02 | -0.17242 | 0 |
| CATCAENFY | HLA-A01:01 | 955.17 | 1.623 | 0.078817 | 0.6 | 0.15364 | 0 |
| FVDENVDRM | HLA-A01:01 | 887.97 | 0.37 | 0.267471 | 0.2 | 0.18947 | 1 |
| HLGVFSSYY | HLA-A01:01 | 711.31 | 0.468 | 0.230519 | 0.2 | -0.13097 | 1 |
| ITSSAELLK | HLA-A11:01 | 18.79 | 0.109 | 0.201403 | 0.2 | -0.10366 | 1 |
| SSSIYSIVK | HLA-A11:01 | 13.64 | 0.127 | 0.365044 | 0.05 | 0.05733 | 0 |
| QALRFYDYK | HLA-A11:01 | 34.77 | 0.849 | -0.097979 | 3 | 0.17556 | 0 |
| KTLKINEVK | HLA-A11:01 | 65.61 | 0.144 | 0.251289 | 0.2 | 0.01153 | 1 |
| ITSFIIERK | HLA-A11:01 | 26.49 | 0.135 | 0.154534 | 0.3 | 0.43372 | 0 |
| AIANRIKFK | HLA-A11:01 | 30.99 | 0.138 | 0.204956 | 0.2 | 0.06827 | 1 |
| ITNQMSIDK | HLA-A11:01 | 48.8 | 0.246 | 0.070952 | 0.6 | -0.32011 | 0 |
| KINEVKTRK | HLA-A11:01 | 79.18 | 0.069 | 0.276622 | 0.1 | -0.00115 | 1 |
| SLSPIEMKK | HLA-A11:01 | 17.93 | 0.023 | 0.18986 | 0.3 | -0.11521 | 1 |
| KMTVVILQK | HLA-A11:01 | 30.02 | 0.172 | 0.140278 | 0.4 | 0.14258 | 1 |
| SIYSIVKIK | HLA-A11:01 | 32.09 | 0.069 | 0.283108 | 0.09 | -0.10345 | 1 |
| QTACCACRK | HLA-A11:01 | 73.54 | 2.384 | 0.309496 | 0.07 | -0.07248 | 0 |
| LSYNSHYSR | HLA-A11:01 | 57.61 | 0.473 | 0.096585 | 0.5 | -0.23814 | 1 |

| | | | | | | | |
|---|---|---|---|---|---|---|---|
| LVANFSQIK | HLA-A11:01 | 28.02 | 0.227 | 0.040298 | 0.8 | -0.05554 | 1 |
| HINTVMEVK | HLA-A11:01 | 96.22 | 0.363 | 0.043777 | 0.7 | 0.02048 | 1 |
| RTMMFFAEH | HLA-A11:01 | 64.68 | 0.852 | 0.076996 | 0.6 | 0.08202 | 1 |
| KANRNLARK | HLA-A11:01 | 126.47 | 0.514 | 0.183548 | 0.3 | 0.0965 | 1 |
| SLFGLSFVR | HLA-A11:01 | 49.75 | 0.518 | 0.126466 | 0.4 | 0.02849 | 1 |
| RVRDFIQMK | HLA-A11:01 | 39.34 | 0.046 | 0.127883 | 0.4 | 0.07804 | 1 |
| TFFLAHGLK | HLA-A11:01 | 370.9 | 2.723 | -0.298934 | 14 | 0.11751 | 0 |

| deepimmuno | mhcflurry_BA | mhcflurry_P | PRIME |
|---|---|---|---|
| 0.57764685 | 11.124151308776 | 0.007375 | 0.186226 |
| 0.6202942 | 14.11473761193 | 0.035375 | 0.218664 |
| 0.75163877 | 25.827487500265 | 0.15725 | 0.165426 |
| 0.830263 | 13.801107492357 | 0.03125 | 0.189427 |
| 0.78717935 | 14.248650007566 | 0.03675 | 0.19647 |
| 0.925292 | 12.372351095531 | 0.016875 | 0.194031 |
| 0.8157916 | 14.908144799284 | 0.041625 | 0.186948 |
| 0.7446863 | 23.488840218842 | 0.134875 | 0.185193 |
| 0.9198542 | 23.313759092692 | 0.132125 | 0.177125 |
| 0.7284804 | 23.373056737771 | 0.132125 | 0 |
| 0.78284633 | 26.219470372838 | 0.15975 | 0.169325 |
| 0.67230046 | 14.023823305792 | 0.033125 | 0.206701 |
| 0.2936206 | 16.124849417643 | 0.054 | 0.192069 |
| 0.79518306 | 19.09961305566 | 0.087625 | 0.194341 |
| 0.6486927 | 14.834287880105 | 0.041625 | 0.183834 |
| 0.21974388 | 17.311998272302 | 0.0695 | 0.192347 |
| 0.7511912 | 30.863959201779 | 0.202 | 0.20391 |
| 0.7671456 | 52.861198035322 | 0.373 | 0.160493 |
| 0.6641836 | 22.380168657242 | 0.122125 | 0.175618 |
| 0.45517206 | 16.159324790747 | 0.054 | 0.205847 |
| 0.6422256 | 19.940328911554 | 0.097 | 0.172907 |
| 0.86775196 | 22.865966216234 | 0.128 | 0.183223 |
| 0.5914246 | 33.134269573105 | 0.223 | 0.171699 |
| 0.801272 | 57.30763461092 | 0.398125 | 0.183394 |
| 0.60707635 | 37.377469770665 | 0.26025 | 0.16699 |
| 0.43357435 | 30.186818621698 | 0.196 | 0.205268 |
| 0.62265426 | 69.073004255867 | 0.4645 | 0.163218 |
| 0.6325165 | 254.86864139858 | 1.002125 | 0.177777 |
| 0.31436318 | 26.135706214331 | 0.000375 | 0.212893 |
| 0.38998526 | 50.728300016666 | 0.042375 | 0.204787 |
| 0.24339715 | 29.435879151756 | 0.00225 | 0.196886 |
| 0.37871394 | 34.361755108306 | 0.008125 | 0.214503 |
| 0.287492 | 101.06084280378 | 0.13925 | 0.212406 |
| 0.23695171 | 50.180289895794 | 0.041125 | 0.197176 |
| 0.18321782 | 424.66292937018 | 0.18625 | 0.17658 |
| 0.38958937 | 49.12488182585 | 0.03925 | 0.213023 |
| 0.61522937 | 176.02506019738 | 0.2465 | 0.201451 |
| 0.4313422 | 37.662202789172 | 0.015375 | 0.181997 |
| 0.44934127 | 85.536175721049 | 0.1125 | 0.194668 |
| 0.48611456 | 52.881240326491 | 0.047875 | 0.199694 |
| 0.40743142 | 142.9717663699 | 0.20425 | 0.187716 |
| 0.37032384 | 74.80535728531 | 0.09325 | 0.19767 |
| 0.2820616 | 167.91698500839 | 0.236 | 0.197812 |
| 0.413391 | 190.2104497628 | 0.104875 | 0.184303 |
| 0.4292046 | 227.92710954986 | 0.30825 | 0.195416 |
| 0.23520824 | 587.94552597755 | 0.616999999999999 | 0.199222 |
| 0.4065638 | 1260.6419846649 | 0.374749999999999 | 0.192327 |
| 0.25980365 | 205.86869586472 | 0.28475 | 0.183613 |
| 0.504692 | 221.13101811976 | 0.302124999999999 | 0.173426 |
| 0.46357292 | 192.16053921279 | 0.267 | 0.18642 |
| 0.31341153 | 116.01611250581 | 0.162125 | 0.181365 |
| 0.53098947 | 183.91634239532 | 0.257625 | 0.186258 |

| | | | |
|---|---|---|---|
| 0.50425655 | 466.92293412367 | 0.532999999999999 | 0.184204 |
| 0.3469938 | 145.71425984689 | 0.207625 | 0.177154 |
| 0.41726232 | 284.65417347797 | 0.367999999999999 | 0.167629 |
| 0.42857555 | 137.17203311583 | 0.193875 | 0.191308 |
| 0.4299041 | 450.43879811043 | 0.520874999999999 | 0.181534 |
| 0.70081514 | 390.20375103233 | 0.17725 | 0.165678 |
| 0.40625614 | 37.478746777189 | 0.009 | 0.191606 |
| 0.26556057 | 29.61423925492 | 0.004125 | 0.203487 |
| 0.40340397 | 289.64574427079 | 0.143625 | 0.165671 |
| 0.56496906 | 185.92502229703 | 0.10325 | 0.165992 |
| 0.44990036 | 25.627058503162 | 0.001875 | 0.187466 |
| 0.27464437 | 50.567433560898 | 0.02 | 0.196196 |
| 0.59653866 | 177.73120797008 | 0.099125 | 0.182886 |
| 0.42737955 | 215.80329608405 | 0.116625 | 0.196076 |
| 0.4255742 | 706.8193028412 | 0.259624999999999 | 0.189817 |
| 0.32530135 | 758.82485307269 | 0.272249999999999 | 0.211075 |
| 0.32538068 | 127.57143639096 | 0.073125 | 0.175985 |
| 0.56712043 | 75.022803237386 | 0.037125 | 0.18432 |
| 0.60504276 | 144.98870205104 | 0.0825 | 0.18654 |
| 0.40528706 | 299.38969425813 | 0.1485 | 0.183884 |
| 0.47778386 | 603.60755506722 | 0.235125 | 0 |
| 0.25645667 | 471.28477036918 | 0.200375 | 0.173896 |
| 0.27098566 | 1204.9718675144 | 0.365624999999999 | 0.160703 |
| 0.40448517 | 90.72209088653 | 0.050875 | 0.18241 |
| 0.35136098 | 258.68435954773 | 0.132 | 0.184697 |
| 0.4339399 | 453.6502965452 | 0.194875 | 0.18891 |
| 0.37788782 | 3052.6391383363 | 0.623749999999999 | 0.173592 |
| 0.20743895 | 716.07475979455 | 0.261499999999999 | 0.185948 |
| 0.41926903 | 143.38561231857 | 0.0815 | 0.19074 |
| 0.5670963 | 1324.801394288 | 0.383124999999999 | 0.176107 |
| 0.22361419 | 10127.446804229 | 1.547625 | 0.164858 |
| 0.25218397 | 1492.5565254185 | 0.409624999999999 | 0.184131 |
| 0.32340282 | 35.98416227662 | 0.01 | 0.200795 |
| 0.21942687 | 1465.7614182063 | 1.06 | 0.183086 |
| 0.3075468 | 193.04721867612 | 0.267 | 0.184123 |
| 0.35086855 | 234.46526043264 | 0.313749999999999 | 0.170838 |
| 0.25650287 | 76.464605086007 | 0.097 | 0.190593 |
| 0.39980733 | 121.06433908365 | 0.169875 | 0.192927 |
| 0.3905387 | 309.04640377954 | 0.393125 | 0.174721 |
| 0.36503214 | 193.66502375083 | 0.270125 | 0.174854 |
| 0.36080515 | 484.6039723727 | 0.542874999999999 | 0.18893 |
| 0.7508924 | 29.241210533714 | 0.05 | 0.177254 |
| 0.5882889 | 35.149604749167 | 0.09725 | 0.203261 |
| 0.839931 | 62.889671812696 | 0.31525 | 0.172644 |
| 0.44005805 | 32.976297551105 | 0.07825 | 0.178267 |
| 0.29303673 | 32.666664525047 | 0.075625 | 0.179739 |
| 0.43898433 | 45.777239461622 | 0.1915 | 0.183036 |
| 0.68718576 | 47.93253807928 | 0.20925 | 0.169394 |
| 0.20847148 | 36.660946791107 | 0.111375 | 0.176649 |
| 0.5301596 | 33.799374883334 | 0.087125 | 0.169477 |
| 0.7422979 | 34.755455708748 | 0.09325 | 0.185412 |
| 0.78236985 | 35.327917779803 | 0.10075 | 0.188694 |
| 0.15599382 | 51.991389966593 | 0.235875 | 0.185631 |
| 0.41065204 | 87.405214173857 | 0.444875 | 0.172559 |

| | | | |
|---|---|---|---|
| 0.4295411 | 44.894235681924 | 0.1845 | 0.174093 |
| 0.6022209 | 68.733373938346 | 0.3465 | 0.177709 |
| 0.18334877 | 105.39772107082 | 0.526375 | 0.180091 |
| 0.5674547 | 66.375067559298 | 0.333375 | 0.174124 |
| 0.93523514 | 47.532225401203 | 0.203875 | 0.183478 |
| 0.39191234 | 30.588265355988 | 0.059375 | 0.18162 |
| 0.52964056 | 549.27255998911 | 1.28375 | 0 |
