## Supplementary Table 3 for "MHC-I binding affinity derived metrics fail to predict tumor specific neoantigen immunogenicity"

**Validation peptides – positive set**

| Author | Paper | Peptide | HLA | CIImm |
|---|---|---|---|---|
| Ehx | https://pubmed.ncbi.nlm.nih.gov/33740418/ | ALDPLLLRI | HLA-A*02:01 | -0.00432 |
| Ehx | https://pubmed.ncbi.nlm.nih.gov/33740418/ | ALPVALPSL | HLA-A*02:01 | -0.04042 |
| Ehx | https://pubmed.ncbi.nlm.nih.gov/33740418/ | SLLSGLLRA | HLA-A*02:01 | -0.12663 |
| Yang | https://pubmed.ncbi.nlm.nih.gov/31011208/ | DKESEEEVS | HLA-C*04:01 | 0.1664 |
| Yang | https://pubmed.ncbi.nlm.nih.gov/31011208/ | DKESEEEVS | HLA-C*12:03 | 0.1664 |
| Huang | https://www.ncbi.nlm.nih.gov/pmc/articles/PMC23 | AVCPWTWLI | HLA-A*11:01 | 0.40404 |
| Yang | https://pubmed.ncbi.nlm.nih.gov/31011208/ | QFIDSSWYL | HLA-A*02:17 | -0.06653 |

| MHCflurry_BA | MHCflurry_P | netMHCpan_R | netMHCpan_BA | Deepimmuno | Deepitope |
|---|---|---|---|---|---|
| 30.78580994879 | 0.202 | 0.023 | 34.56 | 0.86846250295639 | 1 |
| 30.097219147979 | 0.196 | 0.1679 | 73.94 | 0.917633891105652 | 1 |
| 22.265620932586 | 0.122125 | 0.079 | 18.77 | 0.788357377052307 | 1 |
| 24176.537600409 | 18.811875 | 90 | 47731.68 | 0.9283228 | 0 |
| 25540.225082552 | 38.1885 | 100 | 48065.42 | 0.22084391 | 0 |
| 29.745589261505 | 0.05525 | 0.2079 | 36.76 | 0.7719805 | 0 |
| 596.60150256868 | 1.1615 | 0.3948 | 657.37 | 0.81051064 | 1 |

| mixMHCpred_S | mixMHCpred_R | PRIME |
|---|---|---|
| 0.424886 | 0.05 | 0.199219 |
| 0.257277 | 0.2 | 0.177225 |
| 0.431353 | 0.04 | 0.192582 |
| -0.834453 | 90 | 0 |
| -1.299097 | 100 | 0 |
| 0.375697 | 0.04 | 0.225153 |
| 0.022074 | 0.8 | 0.180963 |
