## supplementary table 4 for "MHC-I binding affinity derived metrics fail to predict tumor specific neoantigen immunogenicity"

**Mouse neoantigen**

| Peptide | WT | HLA | BA_mut | BA_WT | %R_mut | %R_WT |
|---|---|---|---|---|---|---|
| FAIFNTEQM | FAIFNTEQR | H-2-Db | 5.21 | 6299.83 | 0.001 | 1.452 |
| VINENYDYL | IINENYDYL | H-2-Db | 63.41 | 65.82 | 0.022 | 0.025 |
| SAYEKLYSL | SAHEKLYSL | H-2-Db | 643.54 | 1492.29 | 0.058 | 0.108 |

Values calculated with netMHCpan 4.1.
*BL: Binding Level.*

| BL_mut | BL_WT |
|---|---|
| SB | WB |
| SB | SB |
| SB | SB |
